# Deep phenomics of human PBMCs using a 31-marker spectral flow cytometry panel

**DOI:** 10.64898/2026.10.31.763346

**Authors:** Soumik Barman, Arsh Patel, Aisling Kelly, Ben Janoschek, Ravi Hingorani, David J. Dowling

## Abstract

Functional cytokines orchestrate leukocyte-mediated innate and adaptive immune responses within the host following infection or immunization. In contrast, lymphocyte-mediated activation-induced markers (AIMs) are predominantly associated with adaptive immune responses. Regulatory T cells (Tregs) primarily function to balance the immune response by preventing excessive inflammation and maintaining self-tolerance. Currently, conventional intracellular cytokine tracking or cytokine-independent AIMs methodology is available for studying antigen-specific lymphocyte or T cell responses in humans. However, this approach faces challenges due to the difficulty in simultaneously deep immunophenotyping the key functional cytokine signatures, AIMs, and nuclear signature of FoxP3^+^ Tregs. Using spectral flow cytometry, we developed a 31-parameter panel to profile simultaneously major immune cell subsets (B cells, NK cells, DCs, monocytes) along with T cells, memory phenotypes, functional cytokines by tracking established Th1 markers IFNγ, TNF and IL-2; Th2 markers IL-4/5/13 and the Th17 marker, IL-17, AIMs and FoxP3^+^ Tregs in human PBMCs in steady state as well as in mitogen stimulated conditions. For AIMs, CD69, CD134 (OX40) and CD137 (4-1BB) are included and validated. We also re-validated our panel to profile antigen specific Th1/2/17 cells or AIMs distribution after peptide stimulation. This optimized multicolor spectral flow panel allows a detailed immune profiling of functional cytokines, AIMs and FoxP3^+^ Tregs in the T cell compartment from human and might be useful for exploratory analysis of how these functional cytokines or AIMs or Tregs shape host immunity after infection or vaccination. Our panel is highly customizable, allowing researchers to tailor it to their specific requirements.

## 1.| Panel Presentation

### 1.1| Purpose and Appropriate Sample Types

This 31-marker spectral flow cytometry panel was meticulously designed and optimized to elucidate the innate and adaptive innate immune profiles in human clinical cohorts. Optimization was carried out using cryopreserved human peripheral blood mononuclear cells (PBMCs) from a healthy adult cohort. Our panel portrays innate live CD45^+^ leukocytes, T cells with memory signature, B cells, monocyte stages, dendritic cells (DCs) and natural killer (NK) cells. To quantify antigen-specific adaptive immune responses, we integrated intracellular staining (ICS) to profile Th1 (IFNγ, TNF, IL-2), Th2 (IL-4/5/13), Th17 (IL-17) cells, and Activation Induced Marker (AIM) assay, which is cytokine-independent. Validation and refinement of AIM-specific markers CD134 (OX-40), CD137 (4-1BB), and CD69 were conducted during ICS or nuclear staining, as this panel was subsequently expanded to profile nuclear FoxP3^+^ regulatory T cells (Treg) signatures in conjunction with AIMs and Th signatures. The panel was validated using various mitogens and peptides to capture antigen-specific T cell responses followed by a kinetics study. This panel is a derivative of the OMIP-XXX-B, which was specifically designed for non-human primates (NHPs) (1). It shares the same objectives as this OMIP. Intentionally, we retained the Th1 signature alongside Tregs as our primary objective to utilize this panel in an opioid use disorder (OUD) cohort (2). Notably, enriched IFN signature (3) and IFN ^+^ FoxP3^+^ cells (4), commonly referred to as “fragile-like Tregs”, are unique characteristics of OUD cohort.

### 1.2| Background

The AIM assay is a functional assay that necessitates the activation of cells through their T cell receptors (TCRs) (5,6). It serves as a robust and highly sensitive alternative for the cytokine-independent detection of antigen-specific T cells (5,6). In parallel, evaluating the cytokine profile of responding T cells may enhance the comprehensive description of the ongoing immune response. During the last quinquennium, hybrid AIM+ICS assays are extensively utilized to evaluate vaccine immunogenicity, particularly in clinical cohorts for emerging or re-emerging zoonotic viruses such as SARS-CoV-2 (7,8), chikungunya (9), monkeypox (10,11), or ICS assays for life-threatening Ebola (12,13) and Nipah viruses (14). Therefore, profiling adaptive immune responses through AIMs (OX-40, 4-1BB, CD69) and intracellular functional cytokines by ICS simultaneously enriched conventional immunophenotyping approaches, despite the often-overlooked cytokine blocker-mediated downregulation of 4-1BB in many OMIPs (Table 1). A few studies have adequately validated the cytokine blocker-mediated downregulation of AIMs (7,15). Limited exploration was conducted to assess the impact of ICS buffer or nuclear stain buffer on profiling adaptive cytokine responses. To address these knowledge gaps in the field of flow cytometry, we developed a unique panel that simultaneously tracks surface AIMs, intracellular functional cytokines, and nuclear proteins (Table 2). This panel has undergone extensive validation using various mitogens with human PBMCs. The NHP-counterpart of this panel has also undergone simultaneous validation (1). For final validation (Figure 1), we chose Concanavalin A (Con A) stimulation for 20 h as AIMs upregulation required an extensive timeframe (16,17). We profiled adaptive cytokine responses as CD69^+^ Th^+^ cells as per OMIPs-5 (18), 52 (19), 67 (20) and 91(21), 20 h post stimulation (Figure 1).

**Figure 1:**
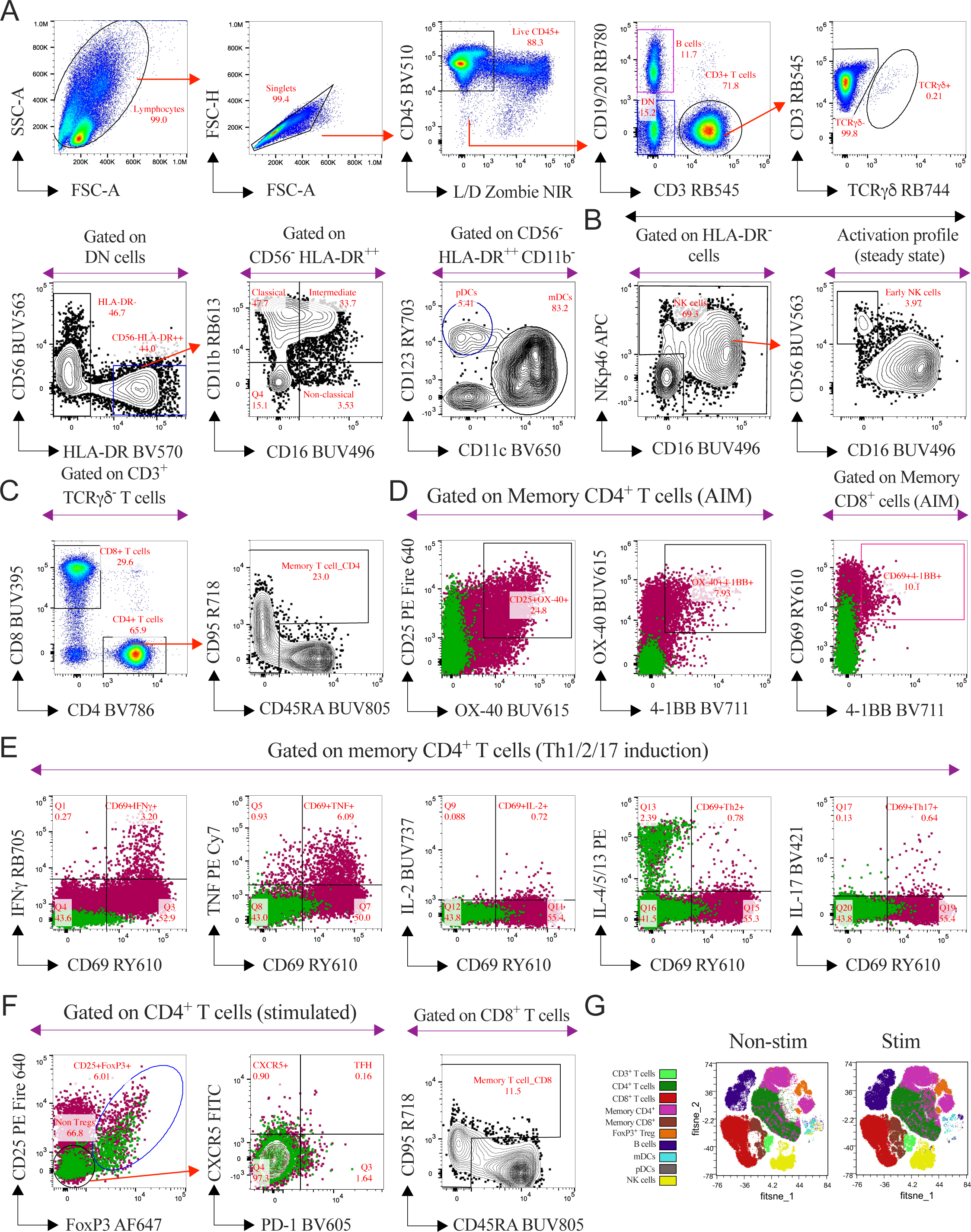
Representative gating strategy to profile AIMs, functional cytokines along with FoxP3^+^ Tregs and FIt-SNE overlays for the visualization of the immune cell populations using 31-parameter OMIP-XXX-A in cryopreserved human PBMCs. (A) Red arrows represent sequential gating for selected populations. After exclusion of cell debris and doublets, gating strategy was shown to profile live CD45^+^ leukocytes, B cells, γδ T cells, monocyte stages (classical, intermediate, non-classical) and DCs. (B) Representative plots to phenotype early NK cells in the NK cell compartment. (C) Gating strategy to portray CD4^+^ and CD8^+^ T cells. Memory compartment was dissected according to the expression of CD45RA and CD95. (D) Representative bivariate plots for profiling AIMs (OX-40 or CD134, 4-1BB or CD137 and CD69) in CD4^+^ or CD8^+^ T cells compartment on rested (clover subsets) or Concanavalin A (Con A) stimulated (cayenne subsets) PBMCs. Activation profiles were visually captured by overlaying clover subsets (steady state) on cayenne (Con A stim, 20 h) cell subsets in the representative bivariate plots. (E) Functional Th1 (IFNγ or TNF or IL-2), Th2 (IL-4, 5 and 13, all together) and Th17 (IL-17) cells were phenotyped in memory CD4^+^ T cells compartment after Con A stimulation for 20 h. In representative bivariate plots, the activation profile (CD69^+^) of Th1/2/17 cell subsets (cayenne) was portrayed by overlaying on rested (clover) subsets. (F) The CD4^+^ T cell compartment was further dissected to profile CD25^+^ FoxP3^+^ Treg subsets and TFH (CXCR5^+^ PD-1^+^) subsets in the non-Treg compartment, either in steady state (clover) or in stimulated (cayenne) condition. (G) 110K events of live CD45^+^ cells from each donor used in concatenated visualization (n=3) with FIt-SNE by the OMIQ platform. The FIt-SNE overlays comparing the topography of cellular landscape in PBMCs in steady state or in Con A stimulated (20 h) condition. For clarity, bivariate plots from the same donor were presented either in steady state or in Con A stimulated conditions.

**Table 1.**
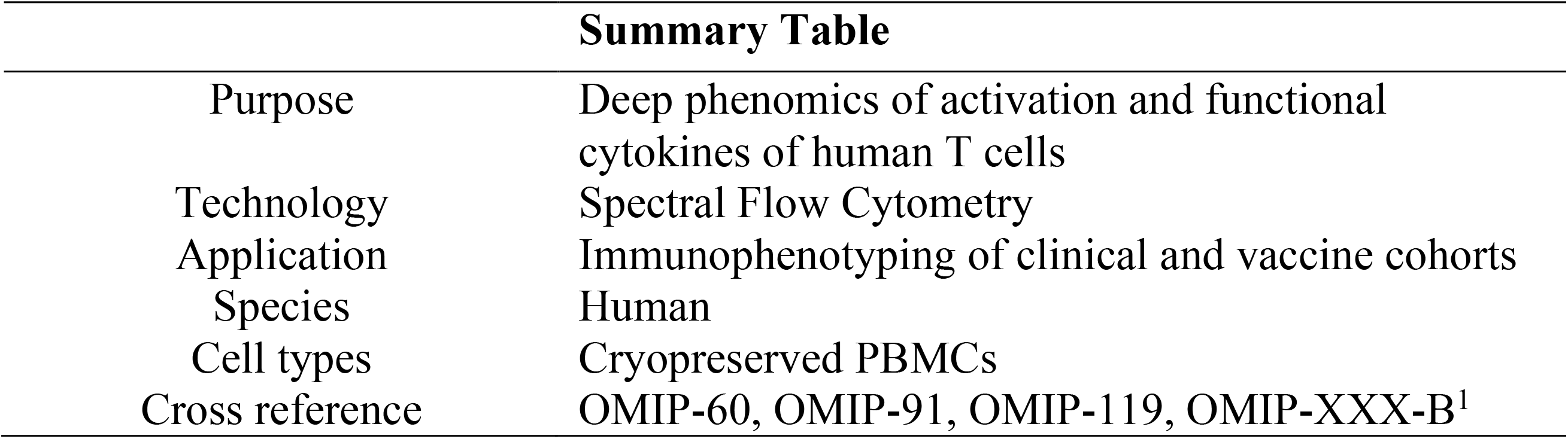
Summary table for the application of OMIP-XXX-A.

| **Summary Table** |  |
|---|---|
| Purpose | Deep phenomics of activation and functional cytokines of human T cells |
| Technology | Spectral Flow Cytometry |
| Application | Immunophenotyping of clinical and vaccine cohorts |
| Species | Human |
| Cell types | Cryopreserved PBMCs |
| Cross reference | OMIP-60, OMIP-91, OMIP-119, OMIP-XXX-B<sup>1</sup> |

**Table 2.**
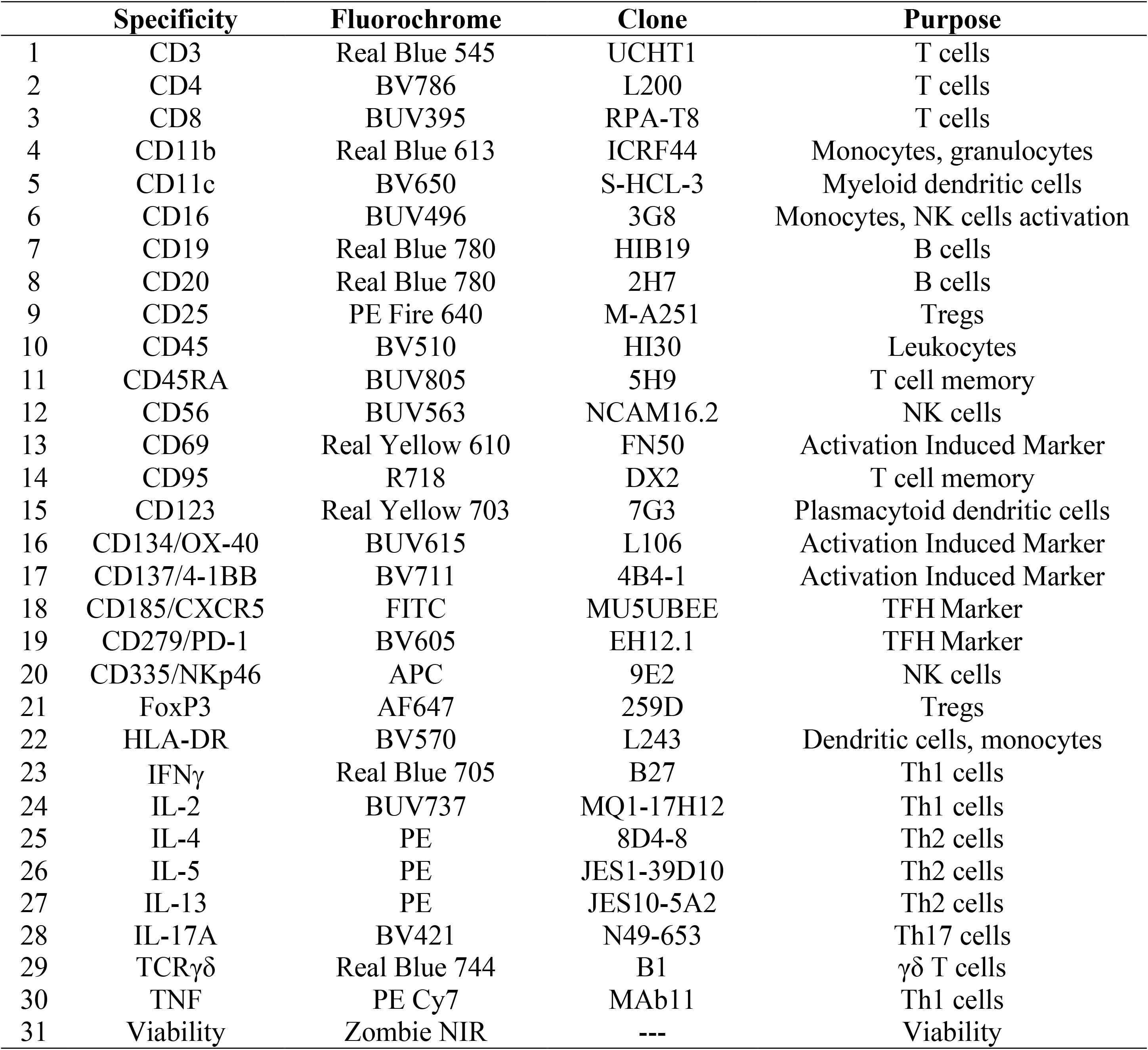
Reagents for OMIP-XXX-A comprising the 31-parameter or the 28-color panel.

|  | **Specificity** | **Fluorochrome** | **Clone** | **Purpose** |
|---|---|---|---|---|
| 1 | CD3 | Real Blue 545 | UCHT1 | T cells |
| 2 | CD4 | BV786 | L200 | T cells |
| 3 | CD8 | BUV395 | RPA-T8 | T cells |
| 4 | CD11b | Real Blue 613 | ICRF44 | Monocytes, granulocytes |
| 5 | CD11c | BV650 | S-HCL-3 | Myeloid dendritic cells |
| 6 | CD16 | BUV496 | 3G8 | Monocytes, NK cells activation |
| 7 | CD19 | Real Blue 780 | HIB19 | B cells |
| 8 | CD20 | Real Blue 780 | 2H7 | B cells |
| 9 | CD25 | PE Fire 640 | M-A251 | Tregs |
| 10 | CD45 | BV510 | HI30 | Leukocytes |
| 11 | CD45RA | BUV805 | 5H9 | T cell memory |
| 12 | CD56 | BUV563 | NCAM16.2 | NK cells |
| 13 | CD69 | Real Yellow 610 | FN50 | Activation Induced Marker |
| 14 | CD95 | R718 | DX2 | T cell memory |
| 15 | CD123 | Real Yellow 703 | 7G3 | Plasmacytoid dendritic cells |
| 16 | CD134/OX-40 | BUV615 | L106 | Activation Induced Marker |
| 17 | CD137/4-1BB | BV711 | 4B4-1 | Activation Induced Marker |
| 18 | CD185/CXCR5 | FITC | MU5UBEE | TFH Marker |
| 19 | CD279/PD-1 | BV605 | EH12.1 | TFH Marker |
| 20 | CD335/NKp46 | APC | 9E2 | NK cells |
| 21 | FoxP3 | AF647 | 259D | Tregs |
| 22 | HLA-DR | BV570 | L243 | Dendritic cells, monocytes |
| 23 | IFNγ | Real Blue 705 | B27 | Th1 cells |
| 24 | IL-2 | BUV737 | MQ1-17H12 | Th1 cells |
| 25 | IL-4 | PE | 8D4-8 | Th2 cells |
| 26 | IL-5 | PE | JES1-39D10 | Th2 cells |
| 27 | IL-13 | PE | JES10-5A2 | Th2 cells |
| 28 | IL-17A | BV421 | N49-653 | Th17 cells |
| 29 | TCRγδ | Real Blue 744 | B1 | γδ T cells |
| 30 | TNF | PE Cy7 | MAb11 | Th1 cells |
| 31 | Viability | Zombie NIR | --- | Viability |

Panel optimization was conducted using a Sony ID7000^TM^ spectral flow cytometer equipped with 5 lasers (355 nm, 405 nm, 488 nm, 561 nm and 637 nm) and 147 fluorescence detectors, as reported in OMIP-111 (22) by our group. The optical layout and fluorescence detector configuration were documented in Table 3. Intentionally, a new generation of fluorochromes is selected (Table 2 & 4) instead of legacy/old dyes to achieve minimal spectral spillovers (Figure 2). Figure 3 also demonstrates minimal spillover spread.

**Figure 2:**
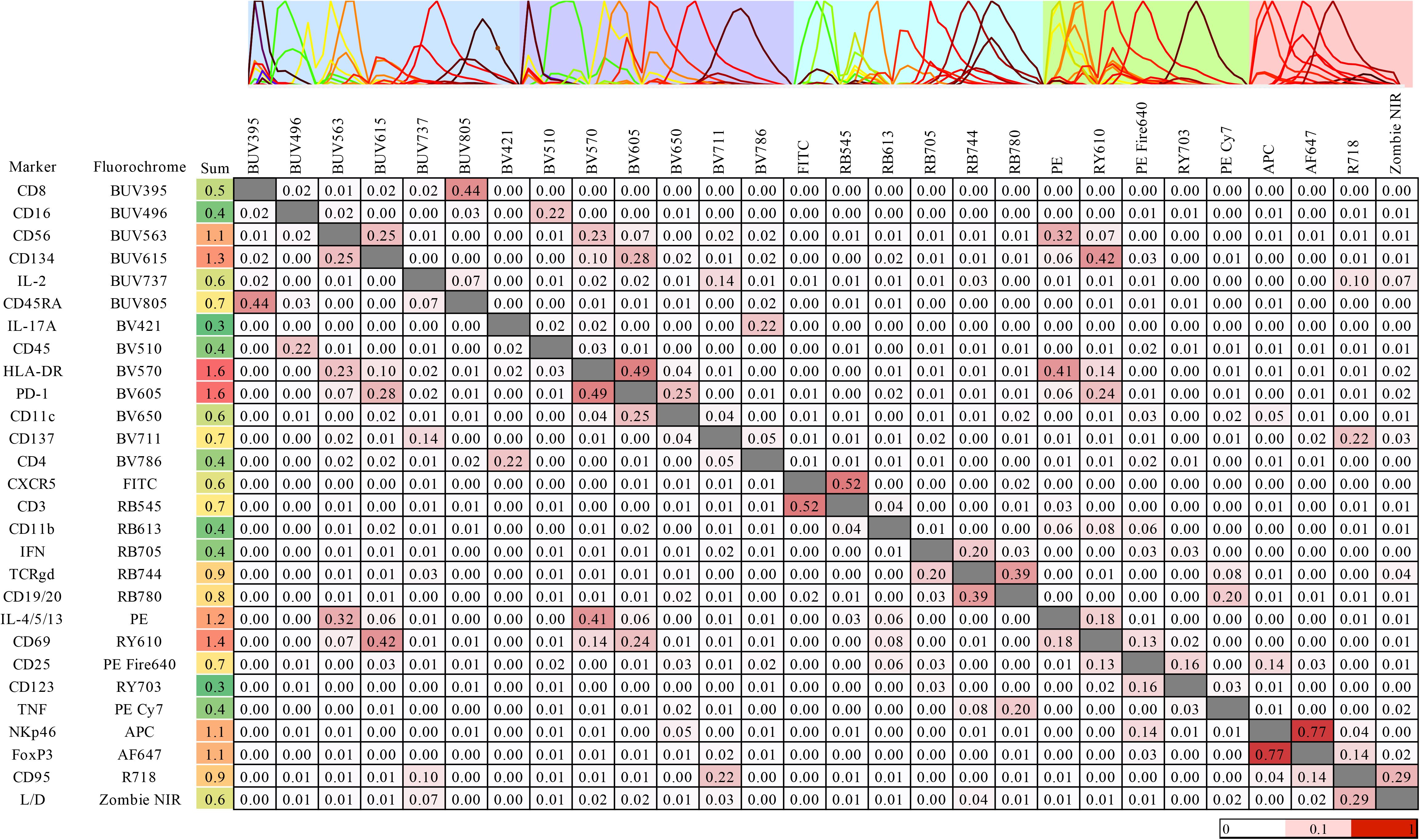
Similarity Index Matrix (SIM) of the optimized 28-color panel on the Sony ID7000^TM^ spectral cell analyzer. The SIM was calculated from in-build spectral references using Panel Development Tool (PDT) developed by Sony Biotechnology Inc. To calculate the similarity score between two fluorophores, normalized fluorescence intensity values of each fluorophore for each detector are plotted and a best fit linear regression line is created. The r^2^value of this linear regression line is the similarity score. The similarity score is on a scale of 0-1 and higher scores indicate a higher similarity between two fluorophores. Higher values in the heatmap are presented in red according to the scale. At the top of the SIM, the spectral distribution was displayed using reference spectra for all the fluorochromes employed in the panel across the instrument’s laser configuration, ensuring precise spectral unmixing. The background colors represent the primary excitation laser line: 355 nm (pale blue), 405 nm (pale purple), 488 nm (ice), 561 nm (lime green) and 637 nm (salmon). Figure was created by Sony Biotechnology’s PDT.

**Figure 3:**
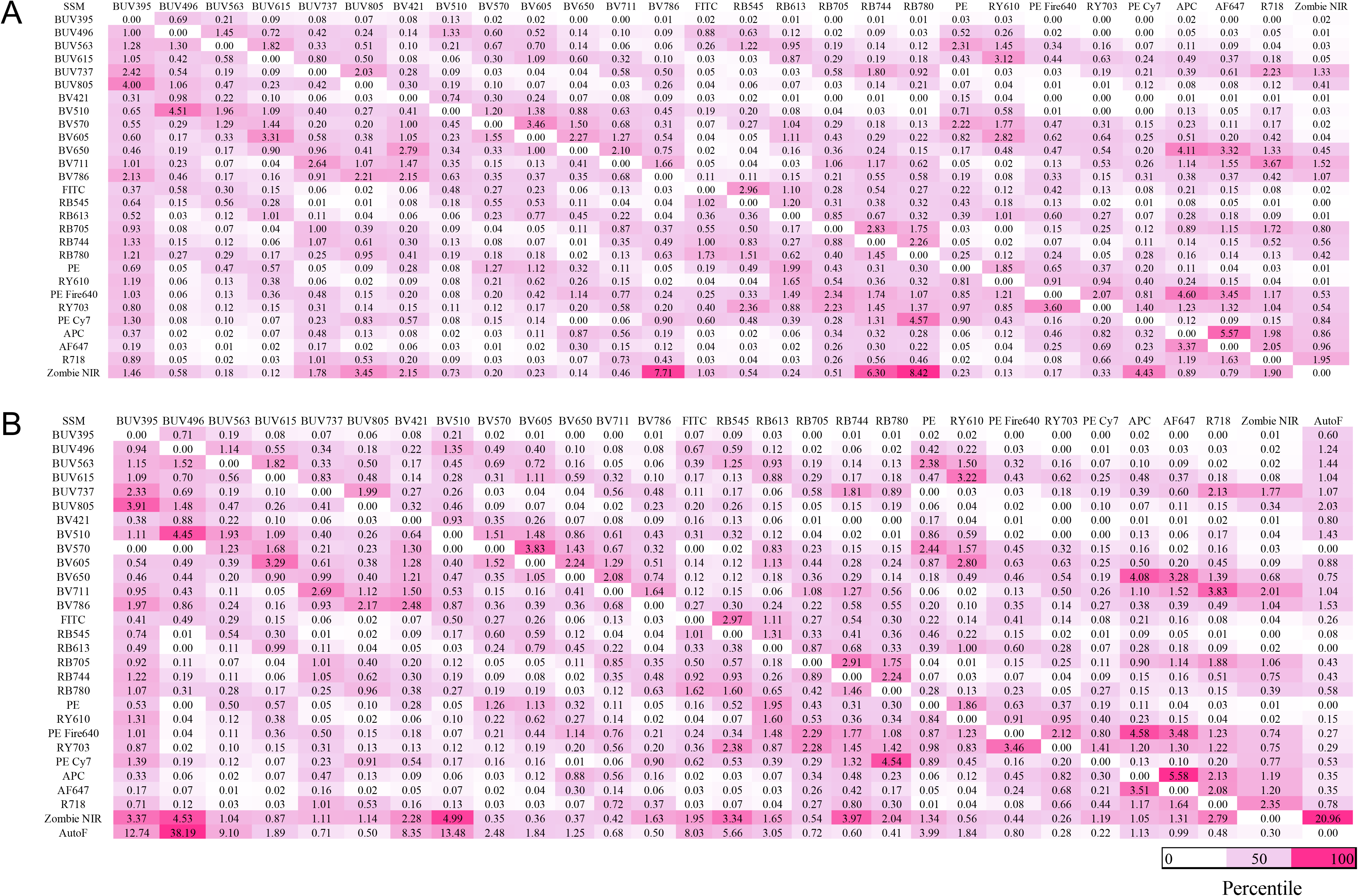
Spillover Spreading Matrix (SSM) of the optimized 28-color panel on the Sony ID7000^TM^ spectral cell analyzer. (A) The SSM was calculated from the single-color controls on antibody capture beads (UltraComp eBeads™ Spectral Unmixing Beads) and utilized to compute the spectral references (via FlowJo 10.10.1 using AutoSpill algorithm). (B) Cellular controls were employed for BV570 and Zombie NIR to mitigate unmixing errors. Additionally, unstained cells were used as an autofluorescence (AutoF) parameter and included in the SSM to subsequently unmix the data presented in this paper. Each cell value provides a quantitative evaluation of the spillover spreading error that a primary parameter’s fluorescence signature (rows/offender fluorophores) generates in the secondary parameter (column, offended fluorophores). Spillover values are presented in light plum (50th percentile) and in strawberry (high) in percentile format.

**Table 3.** Instrumental configuration for the Sony ID7000^TM^. The wavelength range of all spectral detectors is provided. The first two columns provide the channel band and the derived center wavelength (the median of the channel band in nanometers) for each spectral detector on the Sony ID7000^TM^ across all five lasers used for spectral detection. Each spectral detector’s name indicates its position on its laser’s detector array, with the smallest wavelength detector on the left and the largest on the right. Note that the first three detectors for both the UV and Violet lasers have distinct channel bands and center wavelengths, requiring dedicated rows for each laser. The 28 fluorochromes in the panel are also displayed in the channel where they exhibit their maximum fluorescence. The abbreviations for the fluorochromes are as follows: BUV stands for Brilliant UV, BV stands for Brilliant Violet, RB for Real Blue, RY stands for Real Yellow, AF stands for Alexa Fluor and R stands for Red.

| **Detector Array** |  | **Ultraviolet** |  | **Violet** |  | **Blue** |  | **Yellow-Green** |  | **Red** |  |
|---|---|---|---|---|---|---|---|---|---|---|---|
|  |  | **355 nm** | **50 mW** | **405 nm** | **100 mW** | **488 nm** | **150 mW** | **561 nm** | **100 mW** | **637 nm** | **140 mW** |
| Channel Band (nm) | Center Wavelength (nm) | Detector | Fluorochrome | Detector | Fluorochrome | Detector | Fluorochrome | Detector | Fluorochrome | Detector | Fluorochrome |
| 360.5-390 | **375** | **UV1** | **BUV395** | - |  | - |  | - |  | - |  |
| 413.6-438 | **425** | - |  | **V1** | **BV421** | - |  | - |  | - |  |
| 418.5-446 | 430 | UV2 |  | - |  | - |  | - |  | - |  |
| 438-459 | 450 | - |  | V2 |  | - |  | - |  | - |  |
| 446-493.9 | **470** | **UV3** | **BUV496** | - |  | - |  | - |  | - |  |
| 449-480.2 | 465 | - |  | V3 |  | - |  | - |  | - |  |
| 493.9-506.1 | **500** | UV4 |  | V4 |  | **B4** | **FITC** | - |  | - |  |
| 506.1-518.3 | **510** | UV5 |  | **V5** | **BV510** | B5 |  | - |  | - |  |
| 518.3-530.3 | 525 | UV6 |  | V6 |  | B6 |  | - |  | - |  |
| 530.3-542.3 | 535 | UV7 |  | V7 |  | B7 |  | - |  | - |  |
| 542.3-554.2 | **550** | UV8 |  | V8 |  | **B8** | **RB545** | - |  | - |  |
| 554.2-566.1 | **560** | **UV9** | **BUV563** | V9 |  | B9 |  | - |  | - |  |
| 566.1-577.8 | **570** | UV10 |  | **V10** | **BV570** | B10 |  | YG10 |  | - |  |
| 577.8-589.5 | **585** | UV11 |  | V11 |  | B11 |  | **YG11** | **PE** | - |  |
| 589.5-601.1 | 595 | UV12 |  | V12 |  | B12 |  | YG12 |  | - |  |
| 601.1-612.6 | **605** | UV13 |  | **V13** | **BV605** | B13 |  | **YG13** | **RY610** | - |  |
| 612.6-624 | **620** | **UV14** | **BUV615** | V14 |  | **B14** | **RB613** | YG14 |  | - |  |
| 624-635.3 | 630 | UV15 |  | V15 |  | B15 |  | YG15 |  | - |  |
| 635.3-646.6 | **640** | UV16 |  | V16 |  | B16 |  | **YG16** | **PE Fire640** | - |  |
| 646.6-657.7 | **650** | UV17 |  | **V17** | **BV650** | B17 |  | YG17 |  | R17 |  |
| 657.7-668.8 | **665** | UV18 |  | V18 |  | B18 |  | YG18 |  | **R18** | **APC** |
| 668.8-679.8 | **675** | UV19 |  | V19 |  | B19 |  | YG19 |  | **R19** | **AF647** |
| 679.8-690.8 | 685 | UV20 |  | V20 |  | B20 |  | YG20 |  | R20 |  |
| 690.8-701.6 | 695 | UV21 |  | V21 |  | B21 |  | YG21 |  | R21 |  |
| 701.6-712.4 | **705** | UV22 |  | **V22** | **BV711** | **B22** | **RB705** | **YG22** | **RY703** | R22 |  |
| 712.4-723 | **720** | UV23 |  | V23 |  | B23 |  | YG23 |  | **R23** | **R718** |
| 723-733.6 | 730 | UV24 |  | V24 |  | B24 |  | YG24 |  | R24 |  |
| 733.6-744.2 | **740** | **UV25** | **BUV737** | V25 |  | B25 |  | YG25 |  | R25 |  |
| 744.2-756.6 | **750** | UV26 |  | V26 |  | **B26** | **RB744** | YG26 |  | **R26** | **Zombie NIR** |
| 756.6-765 | 760 | UV27 |  | V27 |  | B27 |  | YG27 |  | R27 |  |
| 765-775.2 | 770 | UV28 |  | V28 |  | B28 |  | YG28 |  | R28 |  |
| 775.2-785.4 | **780** | UV29 |  | **V29** | **BV786** | **B29** | **RB780** | **YG29** | **PE-Cy7** | R29 |  |
| 785.4-795.5 | 790 | UV30 |  | V30 |  | B30 |  | YG30 |  | R30 |  |
| 795.5-805.5 | **800** | **UV31** | **BUV805** | V31 |  | B31 |  | YG31 |  | R31 |  |
| 805.5-815.5 | 810 | UV32 |  | V32 |  | B32 |  | YG32 |  | R32 |  |
| 815.5-825.3 | 820 | UV33 |  | V33 |  | B33 |  | YG33 |  | R33 |  |
| 825.3-835.1 | 830 | UV34 |  | V34 |  | B34 |  | YG34 |  | R34 |  |
| 835.1-844.7 | 840 | UV35 |  | V35 |  | B35 |  | YG35 |  | R35 |  |

### 1.3| Results

This panel has been extensively validated using various mitogens such as phorbol myristate acetate (PMA)/Ionomycin, CytoStim, and Con A (Table 5). Con A was selected as a positive control due to its ability to trigger Th1, Th2, and Th17 responses, as well as AIMs (Figure 1).

Figure 1A illustrates the sequential gating strategy employed on a representative donor after nuclear staining to capture live CD45^+^ leukocytes, CD3^+^ T cells, γδ T cells, and B cells using conventional gating strategies for immunoprofiling in steady state. γδ T cells are phenotyped as CD3^+^ TCRgd^+^ according to the OMIPs-69 (23), 84 (24), 99 (25), 102 (26) and 109 (27). The non-B and T cell compartment (CD3^−^CD19/20^−^ or DN) was further dissected to profile monocytes, DCs, and NK cells. Monocytes stages (classical, intermediate, non-classical) were portrayed on CD56^−^ HLA-DR^high/++^ cells according to the expression of CD11b and CD16. The strategy of using CD11b/CD16 expression profile was adopted from the work by the Stanford group (28). Although the use of CD11b instead of the CD14/CD16 combination is uncommon, it is noteworthy that CD14 is ineffective in baboons (1). To ensure consistency, we maintained an identical panel for the non-human counterpart and validated the distribution of CD11b and CD16 across the species (1). Furthermore, we utilized CD56^−^ HLA-DR^high/++^ (non-NK cells) as a parent gate [according to OMIP-109 (27)], followed by gating out granulocyte contamination using CD11b. Subsequently, we dissected the CD56^−^ HLA-DR^high/++^ CD11b^−^ compartment to characterize DCs. Strategy to use CD56^+^ HLA-DR^−^ subset as a parent population for NK cell subpopulation was adopted from OMIPs-58 (29) and 69 (23). Subsequently, the expression of NKp46 and CD16 was employed to characterize NK cells (Figure 1B) according to the OMIP-70 (30). Furthermore, a signature of early NK cells (Figure 1B) was also identified among NK cell subsets by adopting a gating strategy from OMIPs-58 (29) and 69 (23).

After exclusion of γδ T cells from the CD3^+^ T cell compartment, we portrayed CD4^+^ and CD8^+^ T cells and their memory signatures (Figure 1C & 1F). The use of the CD95/CD45RA combination for T cell memory phenotyping is uncommon. However, we justified its application by bypassing either CCR7 or CD28, in conjunction with CD45RA, across human and NHPs (1). Next, we focused on AIMs (Figure 1D) and functional cytokine responses (Figure 1E) in the memory compartments of both CD4^+^ and CD8^+^ T cells after Con A stimulation. The activation profile of the memory CD4^+^ T cell compartment was depicted using the widely used CD25/OX-40 (31,32) and OX-40/4-1BB combinations (7,8,10). The signature of 4-1BB and CD69 (7,10,17) was employed to profile activation stages on memory CD8^+^ T cells. For visualization purposes (Figure 1D-F), subsets in steady state (in clover) were overlaid on their stimulated counterparts (in cayenne). Con A stimulation also successfully induced functional Th1 cells (CD69^+^ IFNγ^+^, CD69^+^ TNF^+^ and CD69^+^ IL-2^+^), Th2 cells (CD69^+^ IL-4/5/13^+^) and Th17 cells (CD69^+^ IL-17^+^) in the memory CD4^+^ T cell compartment (Figure 1E). Functional Th cells were captured as CD69^+^ cytokine^+^ cells. Apart from novel OMIPs (18–21), this strategy was also employed in several recent research studies to profile antigen-specific memory CD4^+^ T cells in vaccine cohorts (33–35). Finally, we profiled Tregs as CD4^+^ CD25^+^ FoxP3^+^ in both steady-state and stimulated conditions, as illustrated in the bivariate overlaid plots in Figure 1F. The non-Treg subset was further phenotyped to identify T Follicular Helper (TFH) cells as CXCR5^+^ PD-1^+^ subsets, as per OMIPs-91 (21) and 99 (25). High dimensionality reduction analysis was performed using the FIt-SNE algorithm by the OMIQ platform to visualize the cellular topography between steady-state and stimulated conditions (Figure 1G). Concatenated visualization (from three healthy donors) suggests the expansion of CD4^+^ T cells (subset in dark green) and memory CD8^+^ T cells (subset in dark brown) upon stimulation.

In vaccine research, CD4^+^ T cell polarization and AIM assay dictate the strength and durability of adaptive immunity (7,9,11). TFH cells directly regulate GC responses and fine-tune long-lasting humoral immunity (36). Tregs are specialized in maintaining immune tolerance (37). Therefore, by employing our distinctive panel, which is first of its kind, in clinical cohorts and vaccine trials, researchers can investigate vaccine-induced AIMs or Th polarization and TFH/Treg induction or alteration. After extensive validation (see **Section 3.2: Panel Optimization**), we finally run our panel using PBMCs from three healthy volunteers and assessed mitogen specific (Figures 1, 11 & 12) and peptide specific T cell response (Figures 11-13). We also validated protocol specific effect of functional cytokine or AIMs detection by comparing ICS and nuclear stain (Figures 11-13). Our primary objective is to utilize this panel for ongoing clinical trials aimed at the development of a vaccine against OUD (38). We speculate that our optimized panel will enable to strategize the pathway to enhance vaccine immunogenicity.

### 1.4| Similarity to Published OMIPs

This is a unique OMIP that is the first of its kind and is available for immunophenotyping of mitogen or antigen-dependent AIM profiles, Th1, Th2, and Th17 cells based on intracellular cytokine signatures along with the nuclear Treg signature (Tables 2 & 4). Conceptually, our OMIP exhibits overlapping characteristics with OMIPs-60 (39), 91 (21), and 119 (40) (Table 1), with a few notable exceptions. OMIP-60 (39) is designed to profile AIMs, Th polarization, and Treg signature. However, the authors prioritized the Treg signature, leading to the omission of validation for protocol-oriented (ICS or nuclear stain) differences that can influence AIMs. The downregulation of 4-1BB by cytokine blockers (see **Section 3.3.2.2: Sensitivity of CXCR5 and AIMs during ICS and Nuclear Stain**) has not been validated or detected in OMIPs-60 (39) and 91 (21). In OMIP-91 (21), authors validated cytokine blocker specific downregulation of CXCR3. In our OMIP, we observed a similar phenomenon in the case of CXCR5 (Figure 10D-I). Consequently, we employed OMIP-91’s strategy (21) to stain chemokine receptors prior to stimulation. In OMIP-119 (40), Th polarization was analyzed using chemokine signatures instead of intracellular cytokines. We adopted the strategy to detect effector cytokines as CD69^+^ cytokine^+^ cells in human PBMCs from the OMIP-91(21). Furthermore, our OMIP has a corresponding NHP counterpart (OMIP-XXX-B; Table 1) designed for and validated in African green monkeys (AGM) and baboons (1). OMIPs-60 (39), 91(21), and 119 (40) are specifically designed for human PBMCs.

## 2.| Technical Information

### 2.1| Instrument’s characteristics, setup and QC

We used a Sony ID7000^TM^ spectral cell analyzer equipped with five lasers (355 nm, 405 nm, 488 nm, 561 nm, and 637 nm) and 147 fluorescence detectors to develop this OMIP. Table 3 presents the optical layout, fluorescence detector configuration, and fluorochrome distribution for the optimized panel. The detector voltage setting was adjusted to achieve optimal resolution for each channel according to Perfetto *et al* (41). The voltages of the FSC and SSC parameters were manually adjusted to distribute the lymphocyte populations. The Dana-Farber Cancer Institute (DFCI) cytometry core conducts daily quality control of the ID7000 using AlignCheck Particles (Sony, Cat# AE700510). Table 4 represents the flow antibodies and their titers used in the 28-color panel designed for Human PBMCs. The titers denote the final dilution factors utilized in the assay.

**Table 4.** Table for flow antibodies used in the optimized 31-parameter or the 28-color panel designed for Human PBMCs. Titer represents the final dilution factors. Each antibody’s volume per test (in 100 μL) was also indicated. The last column indicates the precise staining step where the antibodies were used. Staining of CXCR5 and 4-1BB was not followed in the conventional way due to downregulation and the deviated staining step is marked in magenta. The deviation of markers, clones, fluorochrome, or titer of markers in the human panel compared to its NHP counterpart is highlighted in lime green.

|  | **Specificity** | **Fluorochrome** | **Clone** | **Vendor** | **Identifier** | **Titer** | **(µL/100 µL)** | **Staining Step** |
|---|---|---|---|---|---|---|---|---|
| 1 | CD3 | RB545 | UCHT1 | Waters | 569197 | 1:40 | 2.5 µL | Surface |
| 2 | CD4 | BV786 | L200 | Waters | 563914 | 1:40 | 2.5 µL | Surface |
| 3 | CD8 | BUV395 | RPA-T8 | Waters | 563795 | 1:40 | 2.5 µL | Surface |
| 4 | CD11b | RB613 | ICRF44 | Waters | 571258 | 1:40 | 2.5 µL | Surface |
| 5 | CD11c | BV650 | S-HCL-3 | Waters | 744437 | 1:160 | 0.625 µL | Surface |
| 6 | CD16 | BUV496 | 3G8 | Waters | 612944 | 1:80 | 1.25 µL | Surface |
| 7 | CD19 | RB780 | HIB19 | Waters | 755602 | 1:640 | 0.156 µL | Surface |
| 8 | CD20 | RB780 | 2H7 | Waters | 569120 | 1:20 | 5 µL | Surface |
| 9 | CD25 | PE Fire640 | M-A251 | BioLegend | 356148 | 1:80 | 1.25 µL | Surface |
| 10 | CD45 | BV510 | HI30 | Waters | 563204 | 1:40 | 2.5 µL | Surface |
| 11 | CD45RA | BUV805 | 5H9 | Waters | 742052 | 1:80 | 1.25 µL | Surface |
| 12 | CD56 | BUV563 | NCAM16.2 | Waters | 612928 | 1:160 | 0.625 µL | Surface |
| 13 | CD69 | RY610 | FN50 | Waters | 571167 | 1:80 | 1.25 µL | Surface |
| 14 | CD95 | R718 | DX2 | Waters | 569572 | 1:80 | 1.25 µL | Surface |
| 15 | CD123 | RY703 | 7G3 | Waters | 571445 | 1:40 | 2.5 µL | Surface |
| 16 | CD134/OX-40 | BUV615 | L106 | Waters | 751320 | 1:80 | 1.25 µL | Surface |
| 17 | CD137/4-1BB | BV711 | 4B4-1 | Waters | 569689 | 1:40 | 2.5 µL | Pre-cytokine blockage |
| 18 | CD185/CXCR5 | FITC | MU5UBEE | Thermo Fisher | 11-9185-42 | 1:160 | 0.625 µL | Pre-stimulation |
| 19 | CD279/PD-1 | BV605 | EH12.1 | Waters | 563245 | 1:40 | 2.5 µL | Surface |
| 20 | CD335/NKp46 | APC | 9E2 | BioLegend | 331918 | 1:160 | 0.625 µL | Surface |
| 21 | FoxP3 | AF647 | 259D | BioLegend | 320214 | 1:20 | 5 µL | Nuclear* |
| 22 | HLA-DR | BV570 | L243 | BioLegend | 307638 | 1:40 | 2.5 µL | Surface |
| 23 | IFNγ | RB705 | B27 | Waters | 570247 | 1:20 | 5 µL | Nuclear* |
| 24 | IL-2 | BUV737 | MQ1-17H12 | Waters | 612836 | 1:40 | 2.5 µL | Nuclear |
| 25 | IL-4 | PE | 8D4-8 | Thermo Fisher | 12-7049-42 | 1:40 | 2.5 µL | Nuclear |
| 26 | IL-5 | PE | JES1-39D10 | BioLegend | 500904 | 1:40 | 2.5 µL | Nuclear |
| 27 | IL-13 | PE | JES10-5A2 | BioLegend | 501903 | 1:40 | 2.5 µL | Nuclear |
| 28 | IL-17A | BV421 | N49-653 | Waters | 562933 | 1:40 | 2.5 µL | Nuclear |
| 29 | TCRγδ | RB744 | B1 | Waters | 758180 | 1:20 | 5 µL | Surface |
| 30 | TNF | PE Cy7 | MAb11 | Waters | 557647 | 1:20 | 5 µL | Nuclear |
| 31 | Viability | Zombie NIR | --- | BioLegend | 423106 | 1:1000 | ---- | Post-Fc blockage |

\* The Abs are validated using both ICS and nuclear buffer. However, for the final panel, nuclear buffer was utilized due to the inclusion of FoxP3.

### 2.2| Statement of Ethical Approval, PBMC Isolation and Stimulation

This study involved five healthy human volunteers, comprising four males and one female, with an age range of 21 to 42 years. The optimized panel was finally validated using PBMCs from three out of the five donors. No sex-based analysis was conducted. Blood samples were collected between July and October 2024 and subsequently cryopreserved. Peripheral blood samples were collected after obtaining written informed consent from the volunteers and receiving approval from the Ethics Committee of Boston Children’s Hospital, Boston, Massachusetts (protocol number X07-05-0223). All applicable ethical regulations for human research participants were adhered to.

PBMCs were isolated as described in (42) from fresh whole blood using Ficoll-Paque (Table 5) gradient separation. Cellular viability was assessed using the Cellometer K2 (Nexcelom) instrument with AOPI (Nexcelom) dual staining (Table 5). One mL of a suspension containing 20-25 x 10^6^ cells (per cryovial) was preserved in a solution composed of 90% heat-inactivated FBS (Gibco) and 10% DMSO (Table 5). The samples were stored in liquid nitrogen until their use.

**Table 5.**
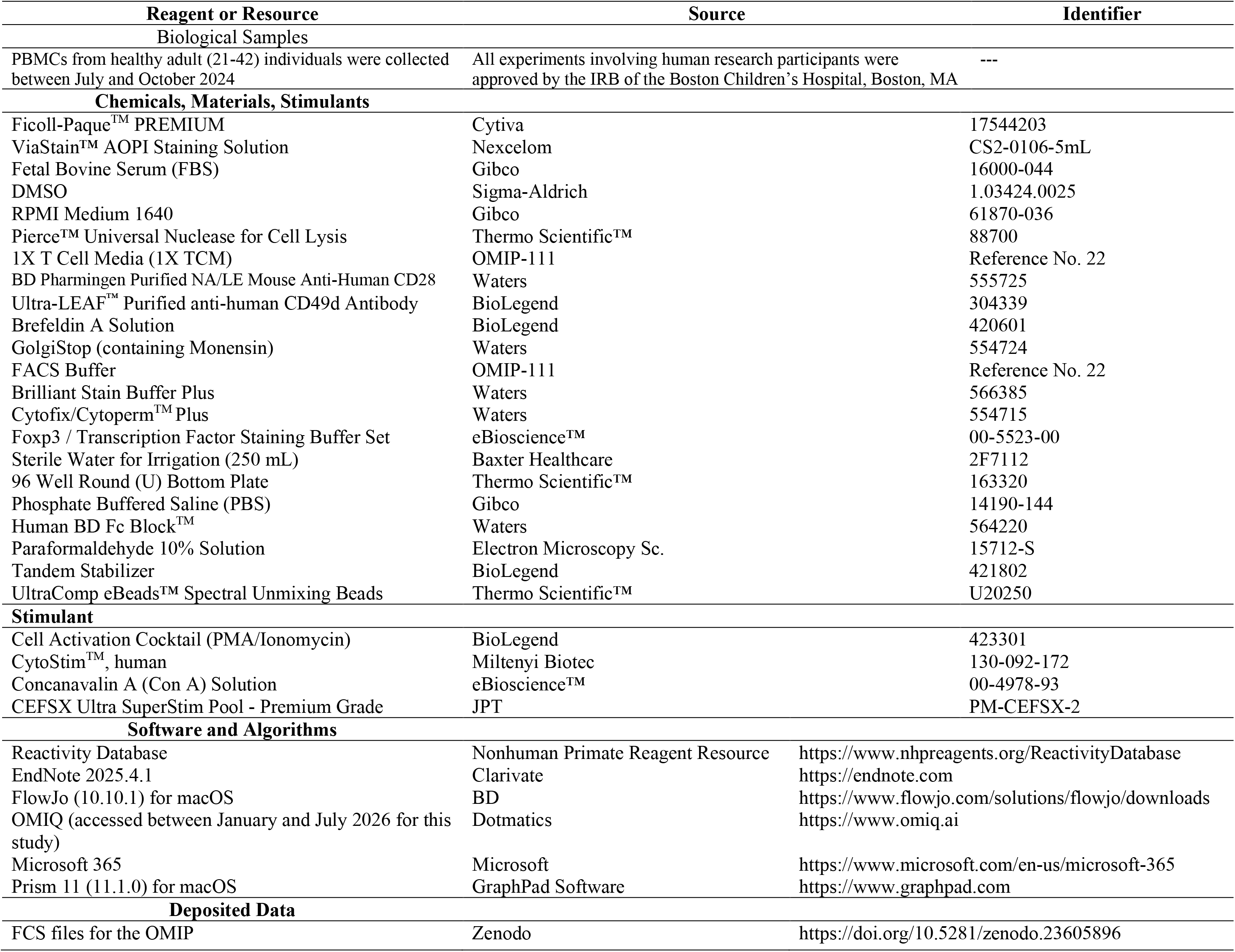
Biological samples, reagents, stimulants and algorithms.

Cryopreserved PBMCs were thawed using a water bath at 37°C. Thawed PBMCs were washed with RPMI 1640 media supplemented with Nuclease (50 Unit/mL) (Table 5). Resting cells in 1X T Cell Media (TCM) (22) were stimulated with mitogens (see **Section 3.3.2.1: Stimulant Optimization**) or peptides (Table 5) and incubated in a humidified incubator at 37°C (in 5% CO_2_). For evaluation of TCR-mediated AIM expression and ICS profiles, cells were stimulated with target peptides in the presence of anti-human CD28/CD49d (1 µg/mL; Table 5). During the last 5 h of stimulation, cytokine production was inhibited using Brefeldin A (1:1000) and Monensin (1:1500) (Table 5). Subsequently, we counted the cells (if necessary) and initiated the staining procedure.

### 2.3| Staining Protocol

#### 2.3.1| Materials and Reagents

Table 5 provides a comprehensive overview of the reagents, buffers, plastic consumables, and software/algorithms utilized in the development of the OMIP. In addition, Table 6 includes reagents that were utilized but ultimately excluded from the optimized panel.

**Table 6.** Commercial reagents were tested but not included in the final optimized panel.

| **Specificity** | **Fluorochrome** | **Clone** | **Vendor** | **Identifier** | **Reason for not using** |
|---|---|---|---|---|---|
| CD3 | BUV496 | SP34-2 | Waters | 573585 | Poor biological separation in NHPs |
| CD56 | BUV496 | R19-760 | Waters | 569468 | Poor biological separation in AGMs |
| CD16 | BUV563 | 3G8 | Waters | 568289 | Better separation in BUV496 |
| TCRγδ | BV650 | B1 | Waters | 564156 | Poor biological separation |
| CD69 | PE CF594 | FN50 | Waters | 562645 | Elevated spillover spreading signatures across multiple channels |
| CD28 | R718 | 28.2 | Waters | 567493 | Changed to CD95 as CD28 was utilized as a co-stimulant during the peptide stimulation |
| CD4 | APC H7 | L200 | Waters | 560837 | This channel was reserved for the NHP panel as we used CD3 in APC Vio770 |
| ArC<sup>TM</sup> reactive beads | -- | -- | Invitrogen™ | A10628 | Unmixing error |
| CytoStim<sup>TM</sup>, NHP | -- | -- | Miltenyi Biotec | 130-094-442 | Unresolved NHP CD3 downregulation under ICS/nuclear staining conditions |

#### 2.3.2| Antibody preparation, Fc blocking and Viability staining

1. During the incubation period, particularly during the final 5 h of cytokine blockade, prepare the cocktail of surface antibodies (Table 4) using FACS buffer (90 µL per test, Table 5) in conjunction with Brilliant Stain Buffer Plus (10 µL per test, Table 5).
2. Prepare 1X Perm/Wash buffer (for ICS) or 1X FoxP3 Permeabilization Buffer (for nuclear staining) by mixing 1 part of stock 10X buffer with 9 parts of irrigated sterile distilled water (Baxter, Table 5). Prepare the cocktail of ICS/nuclear antibodies (Table 4) either using freshly prepared 1X Perm/Wash buffer (90 µL/test) or 1X FoxP3 Permeabilization Buffer (90 µL/test) along with Brilliant Stain Buffer Plus (10 µL/test). For rare surface or functional intracellular/nuclear markers, prepare experimental fluorescence minus one (FMO) or fluorescence minus multiple (FMM) as required.

[**NOTE:** In this OMIP, we validated widely used kits for ICS (Waters, #554715, Table 5) and nuclear staining (eBioscience™, #00-5523-00, Table 5). We conducted ICS and nuclear staining simultaneously to observe the buffer effect, which enables the capture of targeted cytokines or the Treg signature. (see **Section 3.3.2.3: Differences in Cytokine Profile between ICS and Nuclear Stain and Kinetics**)]

1. 3. After preparing the antibody master mix, store it at 4°C in the dark and start the harvesting of rested or stimulated cells for immunophenotyping. Maintain the cellular density (either in steady state or stimulated) up to 1 × 10^6^ cells/well in a 96 well round (U) bottom plate (Table 5) in 200 µL of 1X TCM.
2. 4. Centrifuge (687*g*) the plate contents for 3 min at 18°C. Discard the supernatant by swiftly and cautiously flicking the plate onto the biohazard container containing absorbent soak pads to facilitate rapid absorption of the discarded content.
3. 5. Gently tap the plate to dissolve the cell pellet in the residual media. Subsequently, wash the cells by adding 200 µL of PBS (Gibco, Table 5) and centrifuging at 687*g* for 3 min at 18°C.

[**Note:** For the entire experiment, follow steps 4 and 5 for cell centrifugation, supernatant flicking, and dissolving the cells in the residual volume of the supernatant.]

1. 6. Discard supernatant by flicking and treat the cells with 100 µL of PBS supplemented with Human Fc Block (1.5 µL/test, Table 5) for 10 min at RT. During this incubation period, prepare the viability staining solution (Zombie NIR™, 1:1000 dilution in PBS, Table 4).
2. 7. After incubation, add 100 µL of PBS per well. Repeat step 4 for the wash. Subsequently, stain the cells with the viability staining solution (50 µL/well) for 15 min at RT.
3. 8. After viability staining, add 150 µL of PBS per well. Repeat steps 4 and 5 for a total of two washes.

#### 2.3.3| Surface Antibody Staining

1. Incubate the cells with 100 µL of the surface antibodies cocktail per well for the entire panel run. Ensure all conditions are kept at 4°C in the dark for 30 min.
2. After incubation, add 100 µL of PBS to each well. Execute steps 4 and 5 from **Section 2.3.2**, resulting in a total of two washes at 4°C.

#### 2.3.4| Intracellular or Intranuclear Staining

1. Prepare the cells for ICS by fixing and permeabilizing them. This involves adding 100 µL of BD Cytofix buffer (Table 5) to each well and incubating for 40 min at 4°C in the dark. For nuclear staining, prepare a solution by mixing 1 part of 4X Fix/Perm Concentrate (eBioscience™) with 3 parts of Fix/Perm diluent (eBioscience™). Add freshly prepared 100 µL of this solution to the desired wells for intranuclear staining and incubate the plate for 40 min at 4°C in the dark.
2. Following fixation and permeabilization, add 100 µL of BD 1X Perm/Wash buffer for ICS set. For intranuclear staining set, add 100 µL of 1X FoxP3 Permeabilization Buffer. Repeat steps 4 and 5 from **Section 2.3.2**, resulting in a total of two washes for 4 min (instead of 3 min) at 4°C with 1X required permeabilization buffers.
3. Treat the test wells with either a cocktail (100 µL/well) of targeted intracellular signature or intranuclear signature (Table 4). Incubate all conditions at 4°C for 40 min in the dark.
4. Following the completion of the incubation period, repeat step 2 for washing purposes, resulting in a total of two washes.

#### 2.3.5| Fixation and Storage

1. Next, treat the test wells with 1% paraformaldehyde (200 µL/well) (Table 5). Incubate the plate for 20 min at 4°C. After fixation, wash the wells once with 200 µL PBS/well following centrifugation (687*g*) of the plate contents for 3 min at 4°C.
2. During plate centrifugation, prepare PBS supplemented with Tandem Stabilizer (1:1000, Table 5) and wash the wells once (200 µL/well) with it. Subsequently, preserve the cells in 200 µL of PBS containing the Tandem Stabilizer at 4°C in the dark until acquisition.

#### 2.3.6| Cell Analysis using ID7000^TM^

Stored and fixed cells were analyzed using the Sony ID7000^TM^ spectral cell analyzer within 32 h of staining, unless otherwise specified. Spectral Unmixing Beads (Table 5) were run to define SSM, and test samples were analyzed accordingly. After data acquisition, all fluorochrome combinations (N x N plots) were analyzed using the “unmixing viewer”. Uneven distribution was not adjusted, as it was already addressed with single-color cellular controls (see **Section 3.3.1: Single Color Cellular Controls to Overcome Unmixing Errors**). The new unmixing matrix, combined with beads and cellular controls, was then applied to the entire data set.

## 3.| Panel Design and Optimization

### 3.1| Panel Design

The objective of developing this OMIP was to create a broad-spectral panel that would be suitable for cryopreserved human PBMCs and NHPs, particularly AGM and baboons. This was due to the ongoing vaccine efficacy study utilizing AGM or pertussis pathogenesis using an infant baboon challenge model (1). Consequently, we exercised caution during markers and clone selection. By utilizing the “Reactivity Database” (43) provided by the Nonhuman Primate Reagent Resource (Table 5), we were able to predict the cross-reactivity of the selected clone of targeted markers across three species. We subsequently locked down the clones and proceeded with fluorochrome assignment to the target clones.

#### 3.1.1| Selection of Viability Dye

As an amine-reactive fluorescent dye, we selected Zombie NIR^TM^ due to its unique spectral signature in the red channel. In recent OMIPs-102 (26), 112 (44), and 115 (45), Zombie NIR^TM^ was utilized as a viability dye. OMIP-112 (44) and 115 (45) were designed using Sony ID7000^TM^ spectral cell analyzers, which prompted us to include it in our panel. Furthermore, OMIPs-102 (26) and 115 (45) reported that Zombie NIR^TM^ exhibited lower complexity during the construction of a high-parameter panel.

#### 3.1.2| Fluorochrome Assignment

For the fluorochrome assignment of the selected markers, we utilized PDT developed by Sony Biotechnology Inc. This predictive model (Figure 2) was employed prior to conducting any bench work. In accordance with recent guidelines (46–48), we proceeded to develop a high-parameter panel. During this process, we carefully considered the specific reasons behind assigning fluorochromes to certain markers or targeted clones and documented below based on the primary excitation laser lines.

##### 3.1.2.1| Ultraviolet spectrum (355 nm)

As a bright marker, CD8 was chosen in dimmer fluorochrome BUV395. The combination of CD16 BUV496 and CD56 BUV563 was successfully established in OMIPs-44 (49), 58 (29), 84 (24) and 98 (50). Therefore we implemented this strategy in our OMIPs to track early NK cells using the CD16/CD56 pair after extensive validation (1). The relative brightness of BUV615 and BUV737 is high. Consequently, we decided to retain functional markers CD134 and IL-2 in bright fluorophores as signature of these markers is relative and strictly dependent on the in vitro stimulant. In a high-density marker CD45RA, a dim fluorochrome BUV805 was assigned.

##### 3.1.2.2| Violet spectrum (405 nm)

We utilized IL-17A and CD137 as functional markers, employing bright/moderate fluorochromes BV421 and BV711, respectively. This choice was made since the signature of these markers is relative and entirely dependent on the in vitro stimulant. Due to the high expression of leukocyte marker CD45 and T cell marker CD4, we selected dim fluorochromes BV510 and BV786, respectively. Highly expressed HLA-DR on monocytes and DC was assigned to dim fluorochrome BV570. Since the expression level is low, PD-1 and CD11c were assigned to bright fluorochromes BV605 and BV650, respectively.

##### 3.1.2.3| Blue spectrum (488 nm)

Clone MUSUBEE is used for CXCR5 to track down TFH signature in NHP PBMCs (33) along with human PBMC (51). This clone has already been validated in OMIP-75 (52) for both human and NHPs. As a dim marker, CXCR5 was assigned in moderately bright fluorophore FITC. Except for FITC, for the entire blue spectrum, we utilized the new series of Real Blue dyes. To target the abundant expression of CD3^+^ T cells, dim/moderate fluorophore RB545 was selected. For profiling dim marker CD11b, rare functional cytokine IFNγ, and rare TCRγδ populations, we selected bright fluorochromes RB613, RB705, and RB744, respectively. We intentionally maintained B cell markers CD19 and CD20 in a bright fluorochrome RB780. Notably, employing CD19 (HIB19) RB780 resulted in the inability to identify any B cells in NHP PBMCs (1). Furthermore, CD20 populations are not prevalent in AGM (1). This observation aligns with the trend we observed for γδ T cells in AGM PBMCs (1). In both instances, bright fluorochromes RB780 and RB744 yield exceptional results.

##### 3.1.2.4| Yellow-Green spectrum (561 nm)

To profile functional Th2 cells and the TNF signature, we have selected bright fluorochromes PE and PE Cy7. To identify Th2 responses, we selected IL-4, IL-5, and IL-13 in the same fluorophore PE due to the rarity of Th2^+^ cells within the CD4^+^ T cell compartment. OMIP-56 (53) and OMIP-67 (20) were also followed the same strategy. Of note, in OMIP-56 (53), IL-4 and IL-13 were in APC, while in OMIP-67 (20), IL-4 and IL-5 were assigned to BV421 during the phenotyping of human T cells. Bright fluorochromes RY610 and PE Fire 640 were assigned to AIMs CD69 and CD25 due to their variable signatures. For profiling rare populations of pDCs in cryopreserved PBMCs, we selected bright fluorochrome RY703 for pDC marker CD123.

##### 3.1.2.5| Red spectrum (637 nm)

For dim NK cell marker NKp46, bright fluorochrome APC was chosen. We selected AF647, for FoxP3^+^ T reg signature based on the OMIP-114 (54), as it exhibited the strongest signal compared to other bright fluorochromes (BV421 and PE Cy7). Despite the potential overlap between APC and AF647 (as evidenced by the high similarity index of 0.77 among the scale of 1, Figure 2), we have selected FoxP3 AF647 for our panel after validation (1). As APC (for NK cells) and AF647 (for Tregs) were assigned to distinct cell types, theoretically, this should not result in any unmixing errors. Finally, CD95 was assigned in a moderate fluorochrome R718.

### 3.2| Panel Optimization

#### 3.2.1| Antibody Titrations using combinatorial titration method

A recently developed strategy, “Combinatorial Antibody Titrations” or CAT, which enables the simultaneous titration of three to five antibodies (55), was employed to titrate all the selected antibodies during the panel optimization process (Figure 4-6). In this study, a 2-fold dilution series was employed to achieve most of the antibody’s dilution, encompassing up to 10 distinct dilution points (Figure 5) apart from the viability marker Zombie NIR. Overnight-rested cryopreserved human PBMCs or mitogen-stimulated PBMCs were utilized at an average of 1 million cells per test. Consequently, for each marker’s titration, we employed at least 10 million cells, either in steady-state or stimulated conditions (Figure 5). To mitigate such substantial consumption of AGM and Baboon PBMCs per antibody titration (1), we opted to proceed with the CAT method. Notably, the current panel serves as the counterpart to the NHP backbone panel (1).

**Figure 4:**
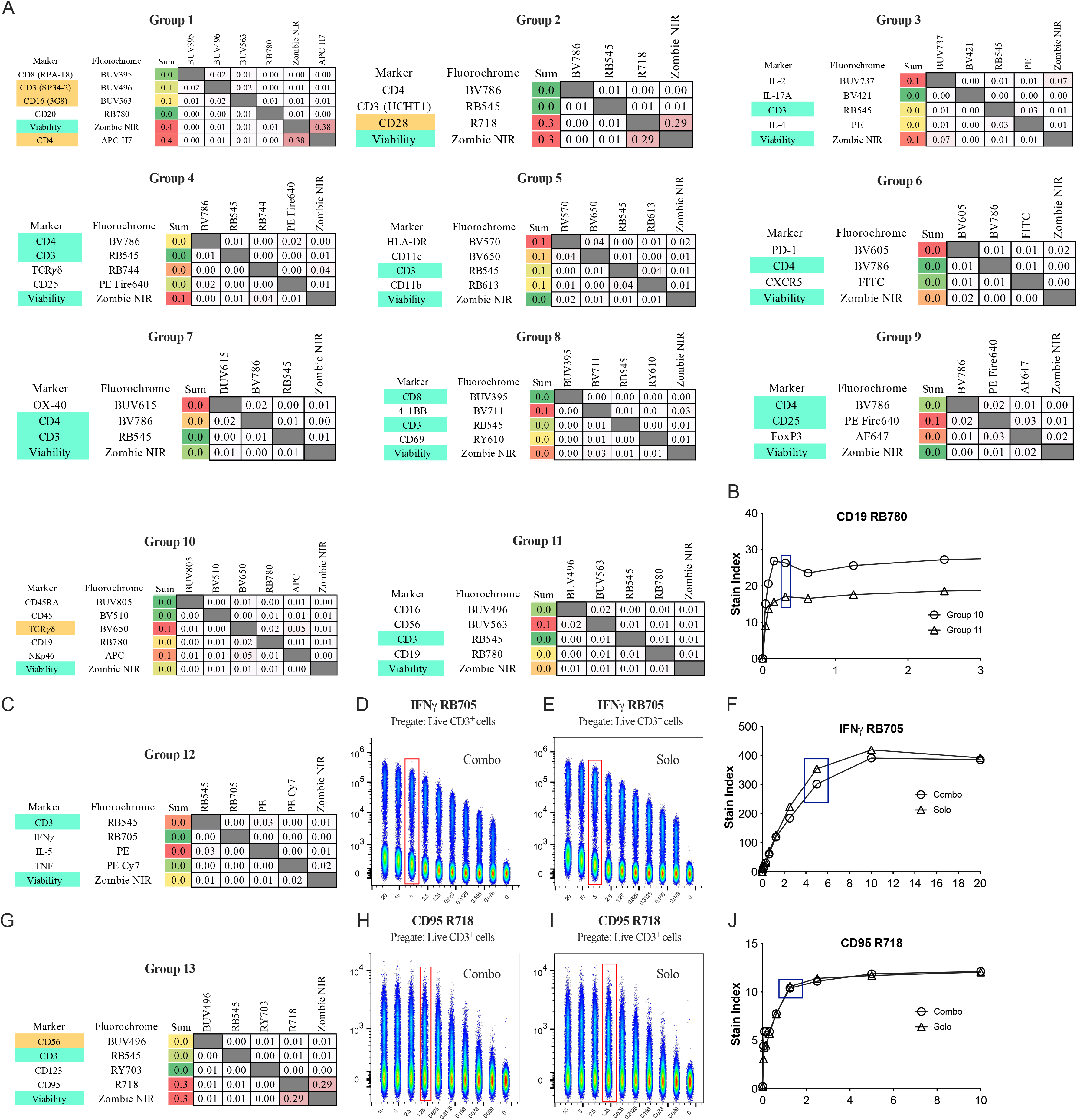
A, C, and G demonstrated the strategy for “combinatorial antibody titrations”. PDT (Sony Biotechnology Inc.) was employed to predict the lowest similarity indices on the SIM, thereby mitigating the risk of unmixing errors. The cut-off value for similarity score was established as less than 0.1 between two fluorochromes. In SIMs, we illustrated the lineage markers in aquamarine. This strategy was devised to titrate rare markers. Markers in orange were utilized during panel development but were not incorporated into the final panel (**Table 6**). Further validation of the “combinatorial antibody titrations” method was conducted by either (B) titrating the same marker in various combinations, or by visually comparing the concatenated representation between combinatorial and traditional methods (D vs E and H vs I). (F, J) Stain Index curve of anonymously selected markers demonstrates the consistency between two distinct titration methods. Chosen titer is indicated by rectangular box in dark blue.

**Figure 5:**
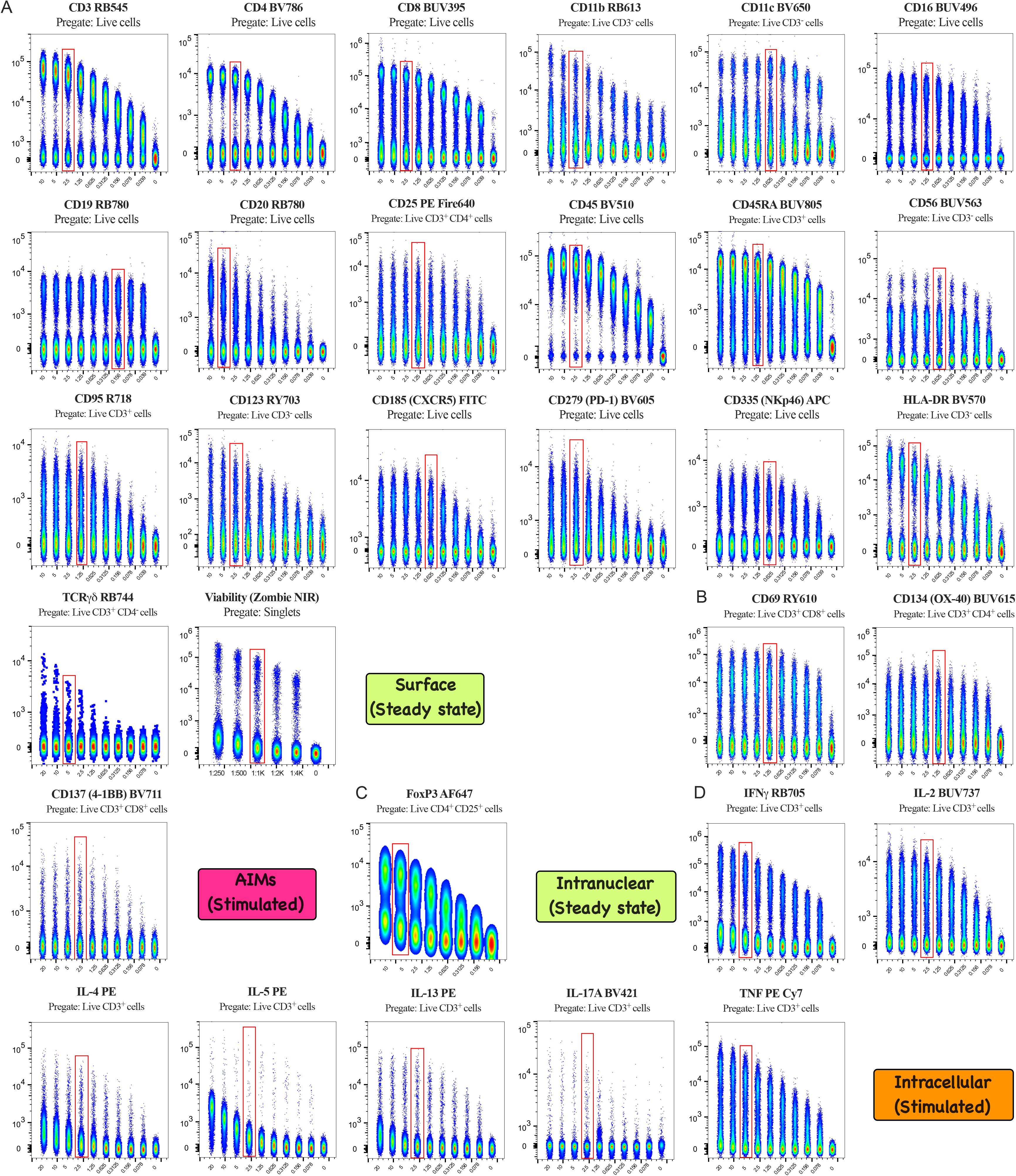
Optimal concentration of antibodies is defined by titrations using overnight rested cryopreserved human PBMCs. Serial fold-dilutions (on the X-axis) of the monoclonal antibodies were started with either 20 µL/100 µL or 10 µL/100 µL, except for L/D Zombie NIR, where the starting dilution was 1:250. All the (A) surface and (C) intranuclear antibodies were titrated using human PBMCs in steady state. For (B) AIMs, cells were stimulated (20 h) with Concanavalin A whereas for (D) intracellular cytokines, cells were stimulated (5 h) with mitogen and titrated thereafter. FCS files were gated to remove debris (FSC/SSC), doublets followed by gating of live cells or indicated subsets (mentioned at the top of each bivariate plots) and concatenated for representation. Red rectangles portray the chosen titer using the Stain Index curve (Figure 6) that provided the distinctive separation between positive and negative populations with minimal background staining. The chosen titration for all the antibodies is also documented as μL/100 μL or as a dilution factor in **Table 4**.

**Figure 6:**
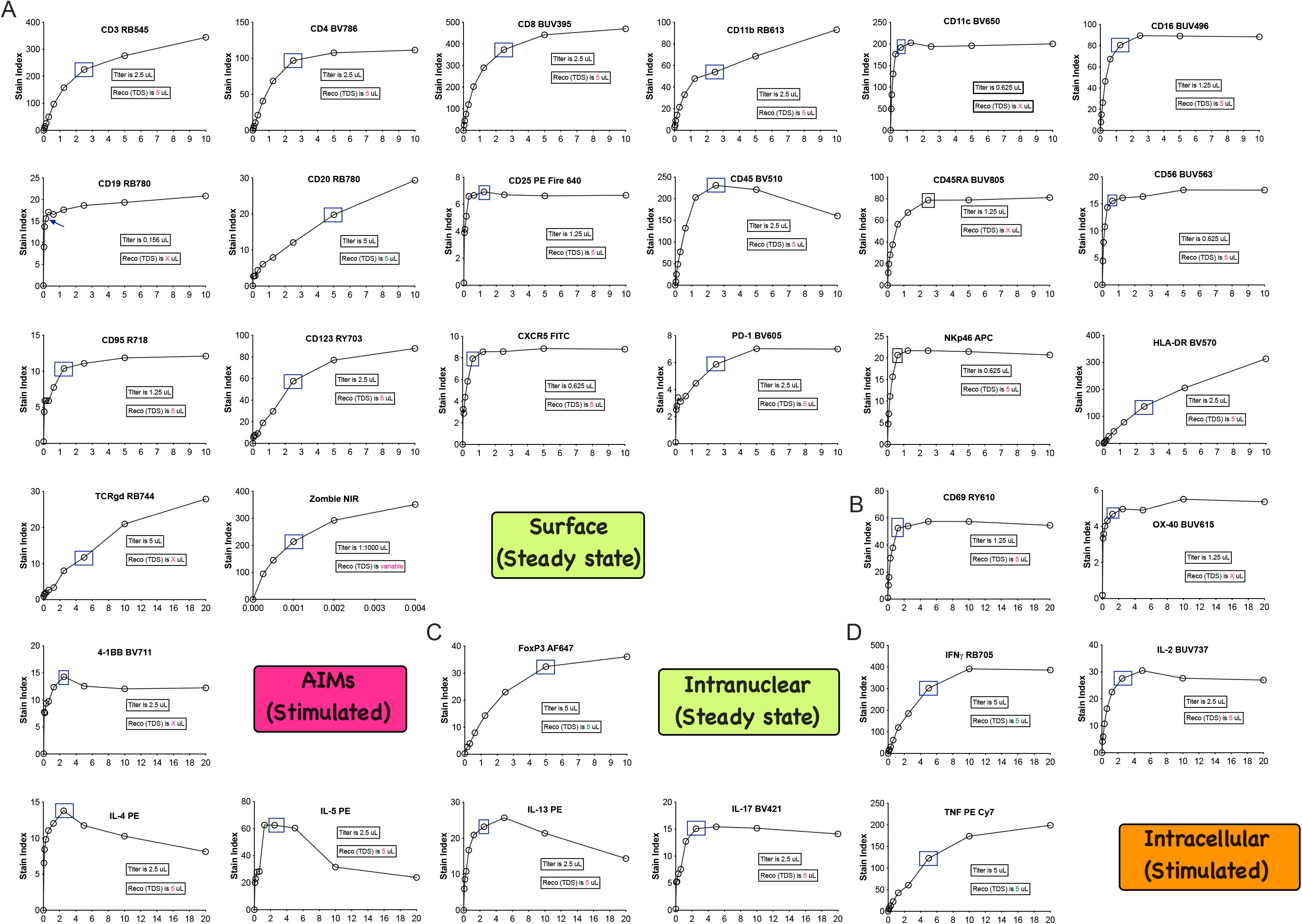
The Stain Index curve of the selected 31 markers for the optimized panel is depicted. A rectangular box in dark blue highlights the chosen titration point where the plateau is observed. The chosen titers are further indicated in each plot. Additionally, the recommended (reco) titration is shown based on the technical data sheet (TDS) of each reagent to provide discrepancies. All the (A) surface, (B) AIMs, (C) intranuclear, and (D) intracellular cytokines were titrated using human PBMCs in either steady-state or stimulated conditions, as illustrated in Figure 5.

Figure 4 showed the strategy to execute the CAT method before any bench work. A total of 13 hypothetical groups were made by assigning the markers in each group with minimal expressional (biological) overlap and with minimal similarity index. The cut off value for similarity index between two (or more) chosen antibodies per group for the CAT was set to ≤0.1 (Figure 4). Viability marker (Zombie NIR) was titrated in solo, and the titrated amount was added thereafter in each group to exclude the dead cells from the analysis. Pre-titrated lineage markers (CD3, CD4, CD8, or CD25) were added as necessary for the selection of cells of interest (highlighted in aquamarine, Figure 4). The pre-gated population utilized for evaluating the titration is depicted in the respective concatenated plots (Figure 5). OX-40 BUV615 was individually titrated (Figure 4, group 7) with the inclusion of lineage markers in the panel. We presented it in Figure 4 as we assessed the combination to avoid spreading errors. Similarly, IL-13 was titrated in solo, but we did not include the SIM in Figure 4 to avoid the potential repetition of either groups 3 or 12.

We also re-validated the effectiveness of CAT to determine whether combined titration influenced the optimal concentration. For re-validation purposes, i) we compared the Stain Index curve of an anonymously selected marker (CD19 RB780) between two distinct groups (Figure 4B); ii) we compared the Stain Index curves of two anonymously selected antibodies (IFNγ RB705 and CD95 R718) that were titrated in combination (combo) or individually (solo) (Figure 4C-J). Similar pattern of Stain Index curves between solo and combo approach (Figure 4F & J) further proved the legitimacy of the CAT method.

The Stain Index was calculated by using FlowJo 10.10.1, using the conventional formula which is Stain Index = [(Med FI of Positive - Med FI of Negative)/(Robust SD of Negative*2)]. After calculation, the stain indices derived from individual markers within each group were plotted to generate a parabolic curve using Prism 11 (Figure 6). In each concatenated bivariate plot (Figure 5), red rectangles indicate the selected titer at the plateau point on the Stain Index curve (Figure 6). These rectangles visually distinguish between positive and negative populations, minimizing background staining. For titration, samples were stained according to **Section 2.3: Staining Protocol**. In this study, all the acquisition related to the titration of antibodies were conducted within a 12-16 h window following the completion of the staining process.

#### 3.2.2| Panel Iterations and FMO Controls

Table 7 outlines the iterations involved in optimizing the panel. Given that the human panel has a corresponding NHP counterpart (1), we aimed to utilize the same clone of antibodies across species. Prior to bench work, we accessed the Nonhuman Primate Reactivity Database (43) to predict reactivity across species and selected the appropriate clones. Subsequently, through iterations 1 to 6 (Table 7), we finalized the clones for phenotyping T cell lineages. Further details are documented in the “Panel Iterations” section of the NHP panel work (1). For each iteration, PBMCs from all species were processed simultaneously. Iterations 23 and 24 were independently assessed to evaluate the performance of PE CF594 and its alternate version, RY610. CD69 PE CF594 did not show potential overlap with BUV615 (Figure 7A) or BV605 (Figure 7B). Of note, CD69 PE CF594 exhibited substantial spillover in RB705 (Figure 7D), RB744 (Figure 7E), PE (Figure 7F), and PE Fire640 (Figure 7G) channels, a phenomenon absent in RY610 fluorochrome. The SSM of iteration 23 and 24 (Figure 7C) further corroborated the potential spillover observed in the bivariate plots. Notably, we incorporated CD3 APC Vio770 in the SSM as a blank channel to assess the spillover mediated by PE CF594, despite its absence in the human panel. It is worth mentioning that CD3 APC Vio770 was utilized in the NHP panel (1).

**Figure 7:**
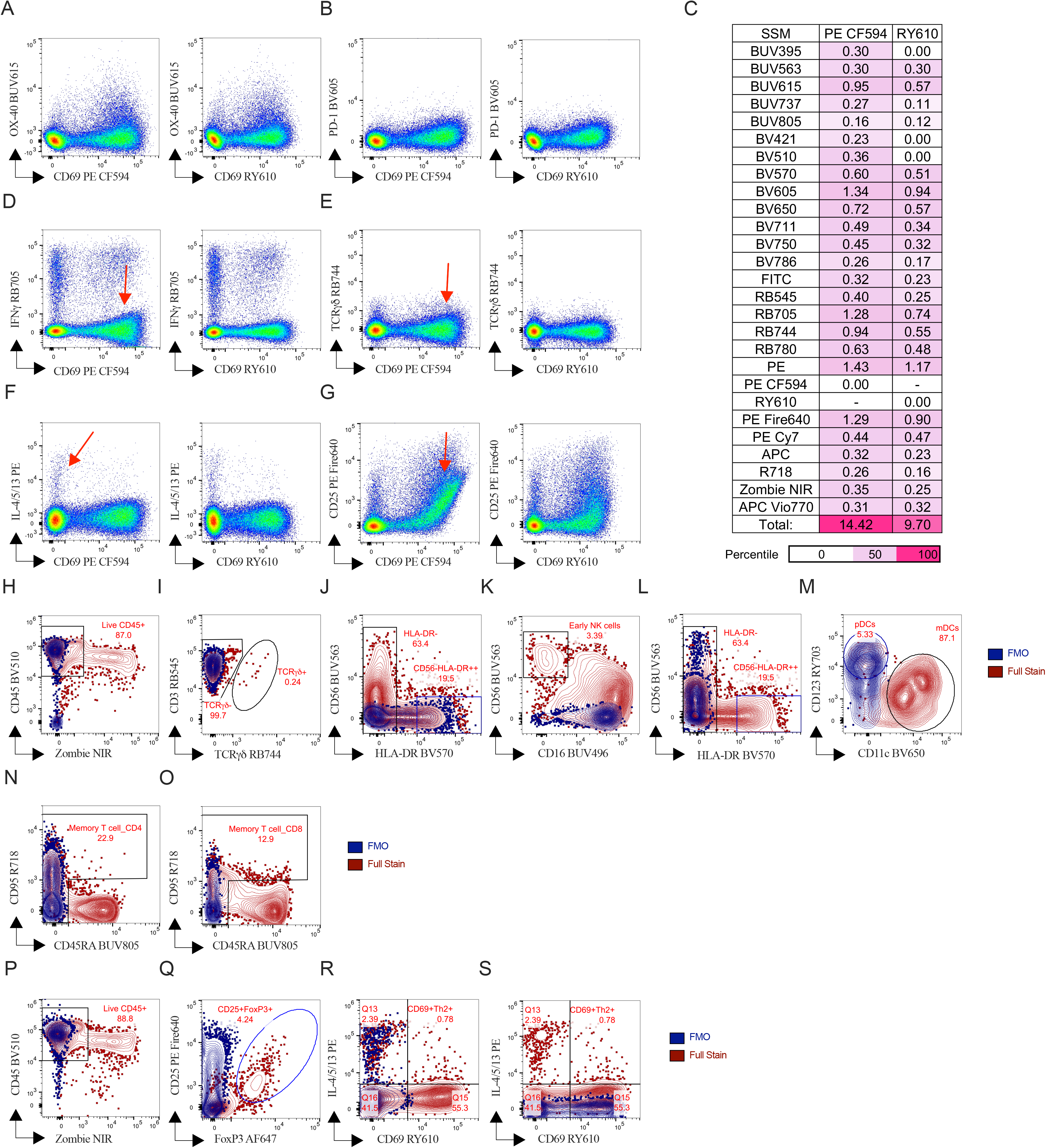
The red arrows indicate potential spillover of CD69 PE CF594 in (D) RB705, (E) RB744, (F) PE, and in (H) PE Fire640. Such spillover can be overcome by using CD69 RY610. RY610 was chosen for the final panel due to its (C) unique minimal spillover spreading signatures in all selected channels used in iterations 23 and 24. Spillover values are presented in (C) light plum (50th percentile) and in strawberry (high) in percentile format. (H-S) Defining gating strategy utilizing FMO controls. To ascertain the accuracy of manually drawn gates, particularly for challenging fluorochromes and rare cell populations, the following FMO plots are presented for (H) live CD45^+^ subset, (I) TCR*γ*δ cells, (J, K) CD56^+^ cells, (L) HLA-DR^high^ cells, (M) CD11c^+^ mDCs and finally (N & O) FMO (CD45RA BUV805) for memory T cells. Cryopreserved PBMCs from reference control were employed in this study for steady-state surface staining without mitogen stimulation. During nuclear staining, FMO plots are presented either for (P) live CD45^+^ subset, (Q) Treg subsets in steady state or (R) CD69^+^ activated T cells, and (S) Th2 cells in Con A stimulated conditions using reference PBMCs. The blue subset (FMO control) was overlaid on the brown subset (full stain) for visual representation.

**Table 7.**
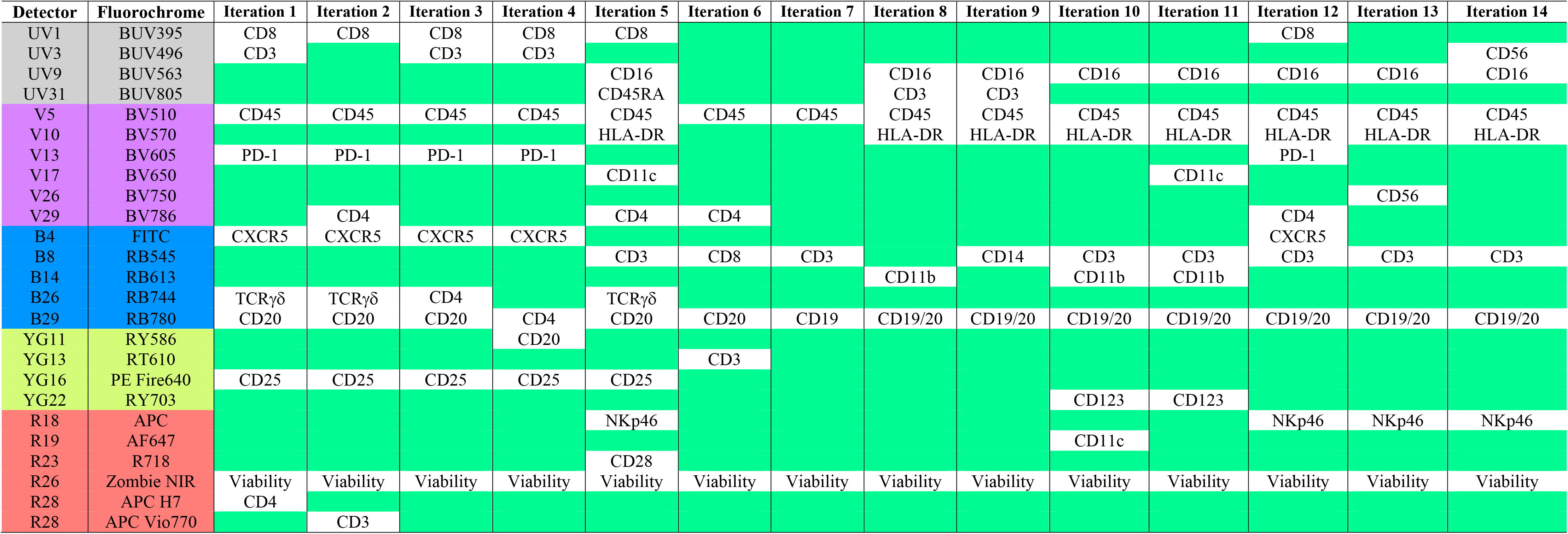
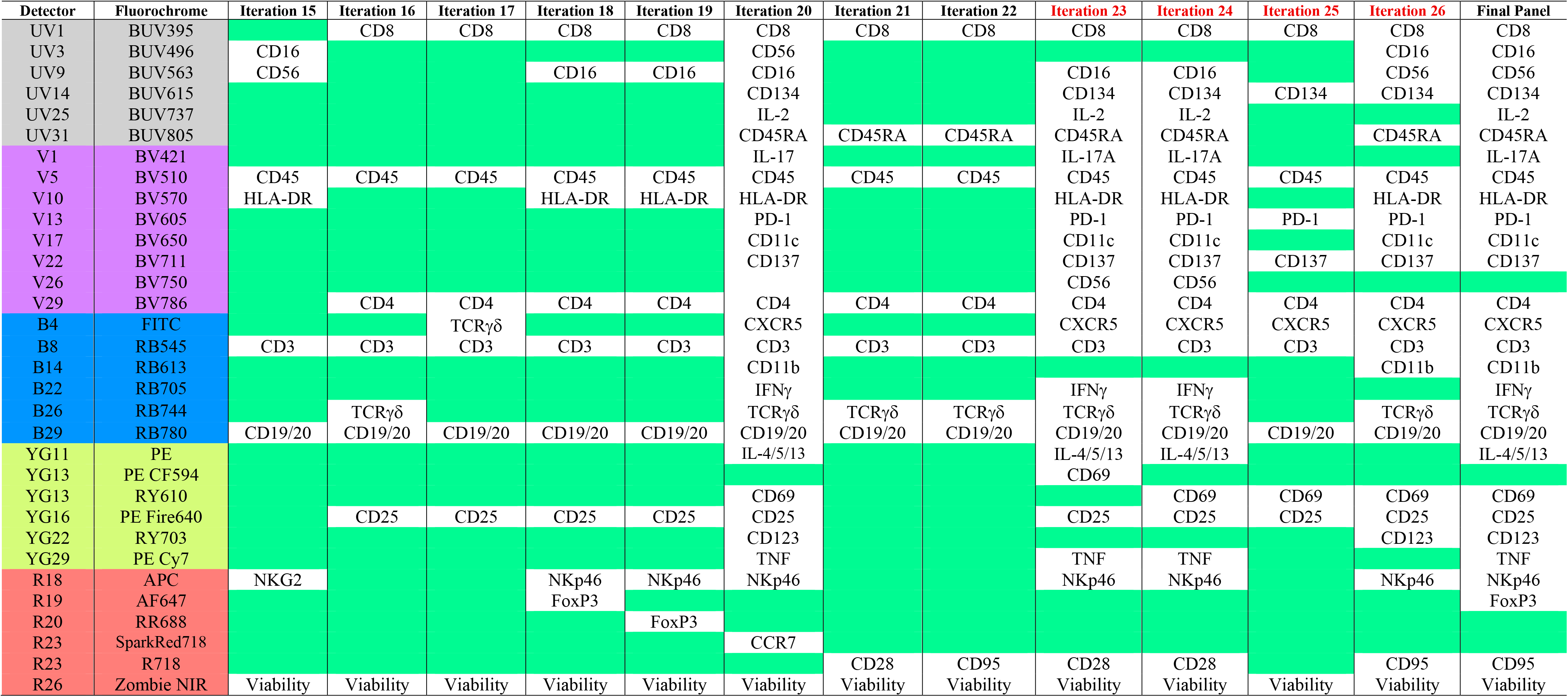
Experiment iterations (1–26) of panel development using human PBMCs. Initially, for each iteration, the NHP version of the panel was simultaneously run using NHP PBMCs. Iterations in red (23–26) were conducted independently.

| **Detector** | **Fluorochrome** | **Iteration 1** | **Iteration 2** | **Iteration 3** | **Iteration 4** | **Iteration 5** | **Iteration 6** | **Iteration 7** | **Iteration 8** | **Iteration 9** | **Iteration 10** | **Iteration 11** | **Iteration 12** | **Iteration 13** | **Iteration 14** |
|---|---|---|---|---|---|---|---|---|---|---|---|---|---|---|---|
| UV1 | BUV395 | CD8 | CD8 | CD8 | CD8 | CD8 |  |  |  |  |  |  | CD8 |  |  |
| UV3 | BUV496 | CD3 |  | CD3 | CD3 |  |  |  |  |  |  |  |  |  | CD56 |
| UV9 | BUV563 |  |  |  |  | CD16 |  |  | CD16 | CD16 | CD16 | CD16 | CD16 | CD16 | CD16 |
| UV31 | BUV805 |  |  |  |  | CD45RA |  |  | CD3 | CD3 |  |  |  |  |  |
| V5 | BV510 | CD45 | CD45 | CD45 | CD45 | CD45 | CD45 | CD45 | CD45 | CD45 | CD45 | CD45 | CD45 | CD45 | CD45 |
| V10 | BV570 |  |  |  |  | HLA-DR |  |  | HLA-DR | HLA-DR | HLA-DR | HLA-DR | HLA-DR | HLA-DR | HLA-DR |
| V13 | BV605 | PD-1 | PD-1 | PD-1 | PD-1 |  |  |  |  |  |  |  | PD-1 |  |  |
| V17 | BV650 |  |  |  |  | CD11c |  |  |  |  |  | CD11c |  |  |  |
| V26 | BV750 |  |  |  |  |  |  |  |  |  |  |  |  | CD56 |  |
| V29 | BV786 |  | CD4 |  |  | CD4 | CD4 |  |  |  |  |  | CD4 |  |  |
| B4 | FITC | CXCR5 | CXCR5 | CXCR5 | CXCR5 |  |  |  |  |  |  |  | CXCR5 |  |  |
| B8 | RB545 |  |  |  |  | CD3 | CD8 | CD3 |  | CD14 | CD3 | CD3 | CD3 | CD3 | CD3 |
| B14 | RB613 |  |  |  |  |  |  |  | CD11b |  | CD11b | CD11b |  |  |  |
| B26 | RB744 | TCRγδ | TCRγδ | CD4 |  | TCRγδ |  |  |  |  |  |  |  |  |  |
| B29 | RB780 | CD20 | CD20 | CD20 | CD4 | CD20 | CD20 | CD19 | CD19/20 | CD19/20 | CD19/20 | CD19/20 | CD19/20 | CD19/20 | CD19/20 |
| YG11 | RY586 |  |  |  | CD20 |  |  |  |  |  |  |  |  |  |  |
| YG13 | RT610 |  |  |  |  |  | CD3 |  |  |  |  |  |  |  |  |
| YG16 | PE Fire640 | CD25 | CD25 | CD25 | CD25 | CD25 |  |  |  |  |  |  |  |  |  |
| YG22 | RY703 |  |  |  |  |  |  |  |  |  | CD123 | CD123 |  |  |  |
| R18 | APC |  |  |  |  | NKp46 |  |  |  |  |  |  | NKp46 | NKp46 | NKp46 |
| R19 | AF647 |  |  |  |  |  |  |  |  |  | CD11c |  |  |  |  |
| R23 | R718 |  |  |  |  | CD28 |  |  |  |  |  |  |  |  |  |
| R26 | Zombie NIR | Viability | Viability | Viability | Viability | Viability | Viability | Viability | Viability | Viability | Viability | Viability | Viability | Viability | Viability |
| R28 | APC H7 | CD4 |  |  |  |  |  |  |  |  |  |  |  |  |  |
| R28 | APC Vio770 |  | CD3 |  |  |  |  |  |  |  |  |  |  |  |  |
| **Detector** | **Fluorochrome** | **Iteration 15** | **Iteration 16** | **Iteration 17** | **Iteration 18** | **Iteration 19** | **Iteration 20** | **Iteration 21** | **Iteration 22** | **Iteration 23** | **Iteration 24** | **Iteration 25** | **Iteration 26** | **Final Panel** |  |
| UV1 | BUV395 |  | CD8 | CD8 | CD8 | CD8 | CD8 | CD8 | CD8 | CD8 | CD8 | CD8 | CD8 | CD8 |  |
| UV3 | BUV496 | CD16 |  |  |  |  | CD56 |  |  |  |  |  | CD16 | CD16 |  |
| UV9 | BUV563 | CD56 |  |  | CD16 | CD16 | CD16 |  |  | CD16 | CD16 |  | CD56 | CD56 |  |
| UV14 | BUV615 |  |  |  |  |  | CD134 |  |  | CD134 | CD134 | CD134 | CD134 | CD134 |  |
| UV25 | BUV737 |  |  |  |  |  | IL-2 |  |  | IL-2 | IL-2 |  |  | IL-2 |  |
| UV31 | BUV805 |  |  |  |  |  | CD45RA | CD45RA | CD45RA | CD45RA | CD45RA |  | CD45RA | CD45RA |  |
| V1 | BV421 |  |  |  |  |  | IL-17 |  |  | IL-17A | IL-17A |  |  | IL-17A |  |
| V5 | BV510 | CD45 | CD45 | CD45 | CD45 | CD45 | CD45 | CD45 | CD45 | CD45 | CD45 | CD45 | CD45 | CD45 |  |
| V10 | BV570 | HLA-DR |  |  | HLA-DR | HLA-DR | HLA-DR |  |  | HLA-DR | HLA-DR |  | HLA-DR | HLA-DR |  |
| V13 | BV605 |  |  |  |  |  | PD-1 |  |  | PD-1 | PD-1 | PD-1 | PD-1 | PD-1 |  |
| V17 | BV650 |  |  |  |  |  | CD11c |  |  | CD11c | CD11c |  | CD11c | CD11c |  |
| V22 | BV711 |  |  |  |  |  | CD137 |  |  | CD137 | CD137 | CD137 | CD137 | CD137 |  |
| V26 | BV750 |  |  |  |  |  |  |  |  | CD56 | CD56 |  |  |  |  |
| V29 | BV786 |  | CD4 | CD4 | CD4 | CD4 | CD4 | CD4 | CD4 | CD4 | CD4 | CD4 | CD4 | CD4 |  |
| B4 | FITC |  |  | TCRγδ |  |  | CXCR5 |  |  | CXCR5 | CXCR5 | CXCR5 | CXCR5 | CXCR5 |  |
| B8 | RB545 | CD3 | CD3 | CD3 | CD3 | CD3 | CD3 | CD3 | CD3 | CD3 | CD3 | CD3 | CD3 | CD3 |  |
| B14 | RB613 |  |  |  |  |  | CD11b |  |  |  |  |  | CD11b | CD11b |  |
| B22 | RB705 |  |  |  |  |  | IFNγ |  |  | IFNγ | IFNγ |  |  | IFNγ |  |
| B26 | RB744 |  | TCRγδ |  |  |  | TCRγδ | TCRγδ | TCRγδ | TCRγδ | TCRγδ |  | TCRγδ | TCRγδ |  |
| B29 | RB780 | CD19/20 | CD19/20 | CD19/20 | CD19/20 | CD19/20 | CD19/20 | CD19/20 | CD19/20 | CD19/20 | CD19/20 | CD19/20 | CD19/20 | CD19/20 |  |
| YG11 | PE |  |  |  |  |  | IL-4/5/13 |  |  | IL-4/5/13 | IL-4/5/13 |  |  | IL-4/5/13 |  |
| YG13 | PE CF594 |  |  |  |  |  |  |  |  | CD69 |  |  |  |  |  |
| YG13 | RY610 |  |  |  |  |  | CD69 |  |  |  | CD69 | CD69 | CD69 | CD69 |  |
| YG16 | PE Fire640 |  | CD25 | CD25 | CD25 | CD25 | CD25 |  |  | CD25 | CD25 | CD25 | CD25 | CD25 |  |
| YG22 | RY703 |  |  |  |  |  | CD123 |  |  |  |  |  | CD123 | CD123 |  |
| YG29 | PE Cy7 |  |  |  |  |  | TNF |  |  | TNF | TNF |  |  | TNF |  |
| R18 | APC | NKG2 |  |  | NKp46 | NKp46 | NKp46 |  |  | NKp46 | NKp46 |  | NKp46 | NKp46 |  |
| R19 | AF647 |  |  |  | FoxP3 |  |  |  |  |  |  |  |  | FoxP3 |  |
| R20 | RR688 |  |  |  |  | FoxP3 |  |  |  |  |  |  |  |  |  |
| R23 | SparkRed718 |  |  |  |  |  | CCR7 |  |  |  |  |  |  |  |  |
| R23 | R718 |  |  |  |  |  |  | CD28 | CD95 | CD28 | CD28 |  | CD95 | CD95 |  |
| R26 | Zombie NIR | Viability | Viability | Viability | Viability | Viability | Viability | Viability | Viability | Viability | Viability | Viability | Viability | Viability |  |

In Figure 7H-S, we delineated the gating strategy for exclusively challenging fluorochromes and rare cell populations using FMO controls. These FMO controls were validated either in steady state (H-O) or in mitogen-stimulated (R & S) conditions to enhance the upregulation of rare cytokine signatures.

### 3.3| Single Color Controls, Other Technical Considerations and Data Analysis

#### 3.3.1| Single Color Cellular Controls to Overcome Unmixing Errors

The combination of antibody-capture beads and cells is reported being useful to achieve the unmixing accuracy (47). Initially, we evaluated our panel using UltraComp eBeads™ Spectral Unmixing Beads (Table 5) along with ArC™ reactive beads (Table 6) for Zombie NIR to compute the SSM. We found unmixing errors especially with Zombie NIR in ultraviolet spectrum (Figure 8A, B), violet spectrum (Figure 8C), blue spectrum (Figure 8D-G), yellow-green spectrum (Figure 8H) and lastly in red spectrum (Figure 8I-K), probably due to ArC^TM^ reactive beads (Table 6). Differences in the autofluorescence spectra between Component A (positive beads) and Component B (negative beads) probably triggers such unmixing errors (26). Using the adjuster of Sony ID7000^TM^ spectral cell analyzer, it is possible to adjust any unmixing errors manually (22,56), but we decided to avoid potential human bias and wanted to rely fully on the Weighted Least Squares Method (WLSM) algorithm. Therefore, we used heat-killed cellular controls (26) for Zombie NIR and as speculated we were able to establish unmixing accuracy (Figure 8A-K, middle column). During further evaluation of each marker in the N x N plots, we noticed potential unmixing error between BV570 and BV421 combo (Figure 8L, red arrows). Further we used cellular controls for BV570 and achieved the accurate unmixing (Figure 8L, bivariate plot in the rightmost section). Unstained samples were treated with the same buffers (PBS, Perm/Wash buffer and nuclear buffer) and used as autofluorescence and included as an independent parameter (as “AF color 1”) during unmixing. We did not extract autofluorescence using the “Autofluorescence Finder” because we did not see any potential issues in our data. Of note, it has been already reported that unnecessary autofluorescence extraction might triggers artifacts (54,57).

**Figure 8:**
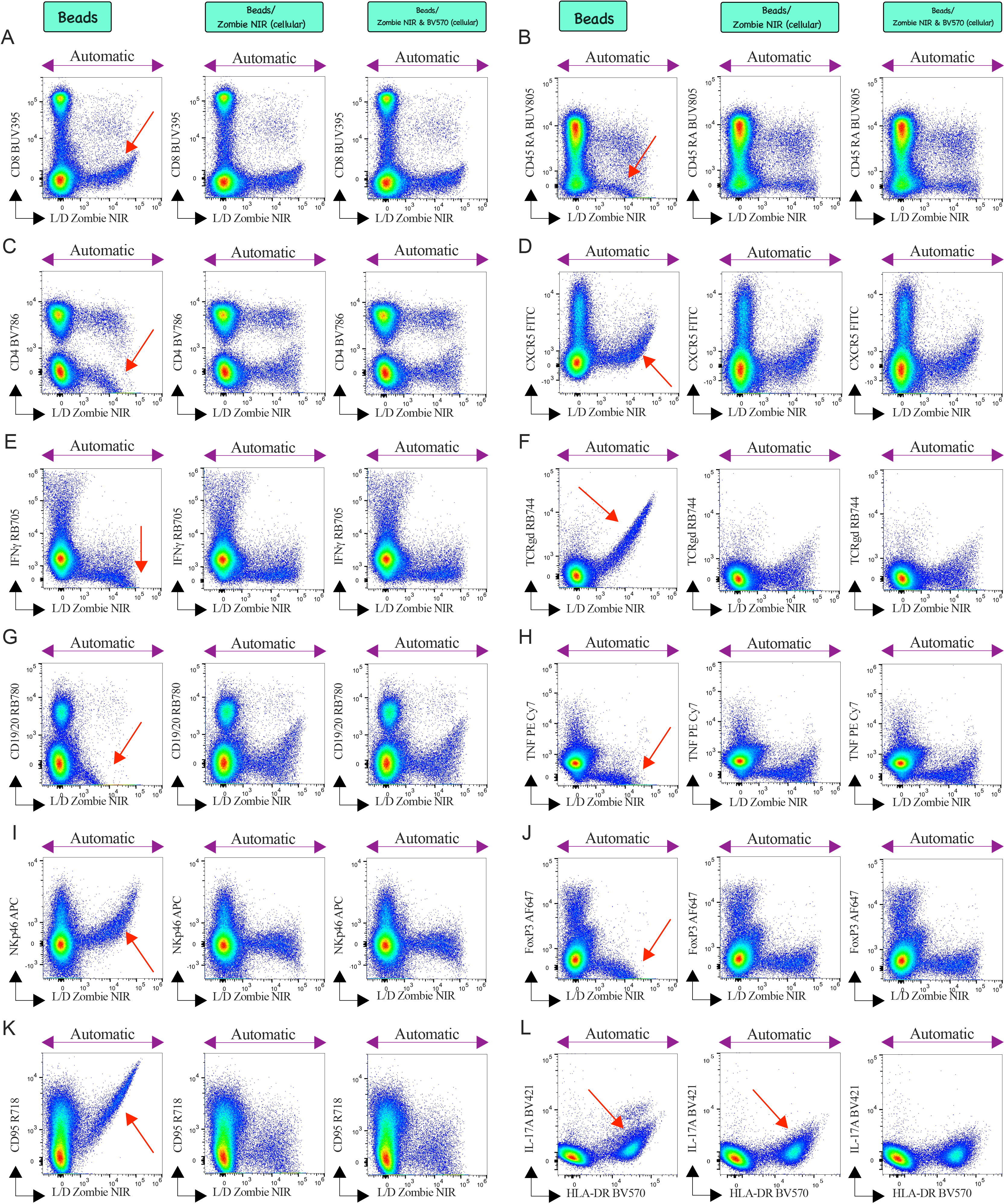
(A-L) Screening of fully stained Con A stimulated (20 h) human PBMCs for detection of unmixing errors (red arrows). N x N matrix was evaluated on singlets for unmixing using UltraComp Spectral eBeads^TM^ along with ArC^TM^ reactive beads (indicated on top as “Beads”). ArC^TM^ reactive beads mediated unmixing errors (red arrows) were shown in (A, B) ultraviolet spectrum, (C, L) violet spectrum, (D-G) blue spectrum, (H) yellow-green spectrum and lastly in (I-K) red spectrum. Such beads mediated unmixing errors can be easily overcome by using cellular controls (denoted as ‘cellular’) especially for Zombie NIR instead of using ArC^TM^ amine reactive compensation beads. Additionally, L) unmixing using Spectral eBeads for HLA-DR BV570 showing potential errors which can be overcome using cellular controls for HLA-DR BV570. ‘Automatic’ denotes no manual adjustment was done using unmixing adjuster in ID7000^TM^.

Step-by-step staining protocol for preparing the reference control for unmixing

1. Plate the optimal concentration (as shown in Figure 5 and Table 4) of the selected antibody conjugates (per well) in a 96-well round (U) bottom plate (Table 5) using FACS buffer (75 µL/test) (Table 5) with the exception of HLA-DR BV570, Zombie NIR and gently mix the contents of each well.
2. After thoroughly vortexing the UltraComp eBeads™ Spectral Unmixing Beads (Table 5), add one drop of the bead in each well. Carefully mix the contents of each well using a multichannel pipette.
3. After 20 min of incubation at RT in dark condition, centrifuge (687*g*, 4 min, 18°C), flick the supernatant, resuspend and store the contents in PBS (200 µL/well) at 4°C (in dark condition) until acquisition.
4. To prepare the reference control for Zombie NIR, generate heat-killed cells by incubating live cells at 65°C for 1 min, followed by 1 min on ice (26), and mix them 1:1 with live cells prior to staining. Use live cellular reference controls for HLA-DR BV570. Unstained live cells served as the negative control for Zombie NIR and HLA-DR BV570 during unmixing. Blocking, surface staining, and fixation were performed as outlined in **Section 2.3.2**.

All acquisitions in this publication were completed within 16 h of the completion of stains with freshly prepared reference controls for each experiment, unless otherwise specified. During acquisition in the ID7000^TM^ spectral cell analyzer, 10,000 reference beads per sample were acquired on every occasion, or 50,000 singlets were acquired for Zombie NIR/HLA-DR. The unmixing process was performed using the WLSM as the unmixing algorithm, and the SSM (Figure 3) was defined.

#### 3.3.2| Other Technical Considerations

##### 3.3.2.1| Stimulant Optimization

One of the most challenging aspects of this panel development was to identify a positive control that can upregulate both AIMs and ICS profiles without compromising the cellular biology of both human and NHP PBMCs. Figure 9 presented a comprehensive validation process for selecting the most suitable stimulant. Mitogens such as PMA and Ionomycin significantly enhance the production of Th1-associated cytokines in both the memory compartment of T cells upon a 5 h stimulation period (Figure 9C). Consequently, we opted to utilize PMA and Ionomycin to determine the titration of markers associated with ICS (Figure 5D). Despite the exceptional induction of Th polarization, we observed a notable downregulation of CD4 expression upon stimulation (Figure 9B). We did not attempt to restore the CD4 downregulation via intracellular staining because certain OMIPs (20,21,39,52) utilized PMA/Ionomycin for short-term (4-6 h) stimulation to induce Th polarization exclusively. Notably, short-term stimulation is insufficient to upregulate AIMs such as OX-40 and 4-1BB (Figure 9C & K), which aligns with previous studies (58,59). Therefore, as an alternative, we used CytoStim^TM^ (Table 5), a mitogen employed as a positive control in a recent study to induce AIMs upon 20 h of stimulation (17). We observed CytoStim^TM^-mediated CD3 downregulation upon stimulation in both human (Figure 9F) and NHP (Figure 9G) PBMCs. Intracellular CD3 staining effectively restores this downregulation in human PBMCs (Figure 9I & L), but not in NHPs (Figure 9H). A 20 h stimulation with CytoStim^TM^ exhibited remarkable efficacy in inducing the AIM signature and Th polarization in human PBMCs, as evidenced by Figure 9N. Of note, extensive stimulation drives slight CD4 downregulation (Figure 9M).

**Figure 9:**
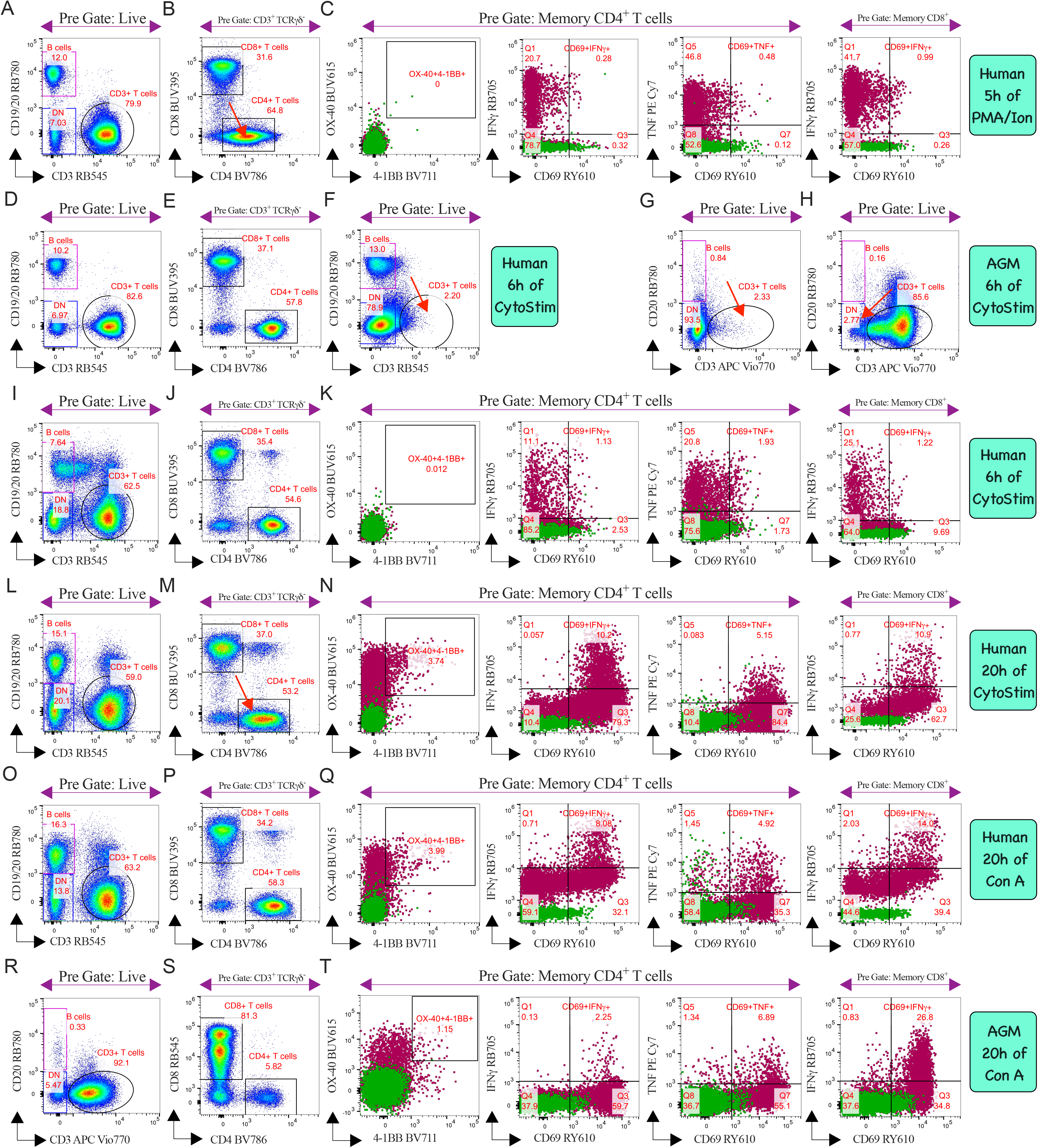
Optimization of stimulants using human PBMCs (A-F & I-Q) and AGM PBMCs (G, H & R-T). All bivariate plots depicted the stimulated conditions, apart from D and E. D and E illustrated the human T cell distribution in the steady state. For TCR-independent stimulation, PMA/Ionomycin (A-C) were evaluated, and a significant downregulation of CD4 expression (B, red arrow) was observed after 5 h of stimulation. CytoStim-mediated CD3 downregulation was also observed in stimulated human PBMCs (F, red arrow) and in stimulated AGM PBMCs (G, red arrow) after 6 h of stimulation. Intracellular staining of CD3 fails to overcome the downregulation observed in AGM (H, red arrow). In humans, CD3 staining in intracellularly restored such downregulation in both 6 h (I) and 20 h (L) of stimulation. Notably, 20 h of CytoStim induces a slight CD4 downregulation (M, red arrow) in human PBMCs. Con A stimulation profiles in humans (O-Q) or in AGM (R-T) were found devoid of such downregulation without alterations of staining protocol and successfully trigger AIMs along with Th polarization (Q, T) in both species. Stimulation profiles (C, K, N, Q, and T) were visually assessed by overlaying clover (steady-state) subsets on cayenne subsets (mitogen stimulation) in the representative bivariate plots. Stimulant conditions and species are listed at the far right of each row.

Ultimately, we validated Con A, which successfully met all the requirements (inducing both AIMs and Th polarization simultaneously) and demonstrated no downregulation of immune cell subsets in both human (Figures 9O-9Q) and NHPs (Figures 9R-9T). Additionally, Con A does not necessitate the bypassing of traditional staining methods. Consequently, we decided to utilize Con A as a positive control for the entire study, extending up to 20 h of stimulation. Notably, AIMs-related markers were also titrated using Con A-stimulated PBMCs (Figure 5B).

We further validated our panel by utilizing peptide stimulation to track TCR-dependent AIMs and ICS profiles. This was achieved using CEFSX Ultra SuperStim peptide pull (Table 5), a combination of 202 peptides and an extended version of the CEFX SuperStim pull. The CEFX SuperStim pull has been extensively utilized to profile AIMs (60) or Th polarization (61) in clinical cohorts. Therefore, we hypothesized that the extensive epitope coverage of CEFSX Ultra SuperStim peptide pull would undoubtedly elicit adaptive responses in healthy individual’s PBMCs. Initially, we investigated whether the presence of any protocol-specific (surface, ICS, or nuclear stain) downregulation of CD3, CD4, and CD8 expression on 20 h of stimulated PBMCs with CEFSX Ultra SuperStim peptide (Figure 10A-C). We did not observe any downregulation upon peptide stimulation, regardless of whether we performed surface staining only (Figure 10A), surface staining followed by ICS (Figure 10B), or nuclear staining (Figure 10C). Subsequently, the detailed process of further fine-tuning and comprehensive validation of our panel is discussed in later sections using PBMCs from three healthy adult human donors.

**Figure 10:**
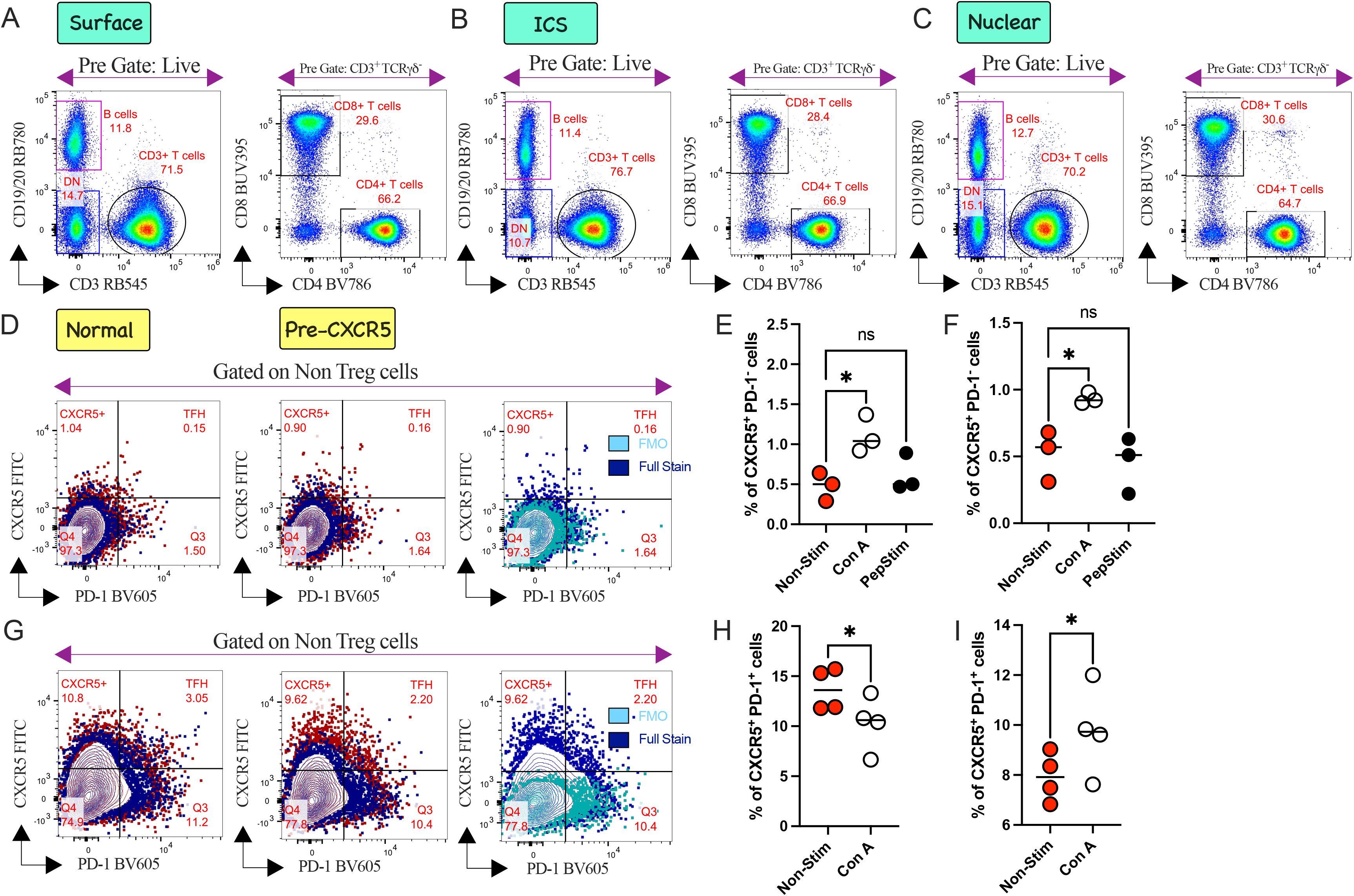
(A-C) Validation of T cell markers CD3, CD4, and CD8 upon CEFSX Ultra SuperStim peptide stimulation for 20 h. Bivariate plots illustrate the distribution of T cell markers after (A) surface staining, (B) ICS and (C) nuclear staining. For (A) surface staining, intracellular markers were not included in the cocktail. The distinct steps of immunophenotyping are also depicted at the top. Assessment of CXCR5 expression upon Con A stimulation using either human PBMCs (D-F) or AGM PBMCs (G-I) after nuclear staining. Each column represents the distinct steps of immunophenotyping as indicated at the top. CXCR5 FITC was added either during surface staining (denoted as Normal) or added 15 minutes before the addition of the stimulant (termed as Pre-CXCR5). Activation profiles were visually captured by overlaying blue (steady state) subsets on the brown subsets (Con A stimulation, 20 h) in the representative bivariate plots (from an individual donor). The sky subset (FMO control) was overlaid on the blue subset (full stain upon stimulation) for visual representation and to justify the gating strategy. Graphical presentation of the CXCR5 profile during normal staining (E, H) or pre-CXCR5 (F, I) after stimulation in human (D-F) and AGM (G-I) respectively, illustrates the magnitude of the stimulant-specific variation of CXCR5 profile in CD4^+^ T cells compartment. Statistical comparison was performed using either two-tailed paired t-test or Wilcoxon test corrected for multiple comparisons; *P <0.05, ns denoted non-significant. Each circle represents a single participant (n=3-4 per group).

##### 3.3.2.2| Sensitivity of CXCR5 and AIMs during ICS and Nuclear Stain

We finally validated our panel using PBMCs from three healthy human donors. Rested cryopreserved PBMCs were either stimulated with Con A as a positive control or CEFSX Ultra SuperStim peptide to portray the adaptive immune response. First, we focused on the specific downregulation of CXCR5 receptors in both human (Figure 10D-F) and AGMs (Figure 10G-I). Stimulant-mediated downregulation of CXCR5 has been previously reported in Rhesus Macaques (19). To address this issue, we added CXCR5 FITC antibody 15 minutes prior to the addition of the stimulus and incubated at 37°C, which we referred to as “Pre-CXCR5” (Figures 10D & G, plots in the middle column; 10F, 10I). This strategy was employed in several studies involving either human (6,8,11,62) or NHP cohorts (63,64). In human, we were unable to detect Con A-specific or peptide-specific CXCR5 downregulation during normal stain (Figure 10E). We successfully captured Con A-specific CXCR5 upregulation whether we added CXCR5 FITC during the surface stain (Figure 10E) or prior to stimulation (Figure 10F). In AGM, we observed Con A-mediated significant downregulation of CXCR5 (Figure 10G, H). Such downregulation could be easily restored using the “Pre-CXCR5” strategy in AGM (Figure 10I). Therefore, we decided to follow this strategy for both human and NHP panels.

Further we examined cytokine blocker (Brefeldin A and Monensin) specific downregulation of 4-1BB (Figure 11A, B, E, F, I, K, M & O) as reported by other studies (7,15). We bypassed such downregulation by adding 4-1BB before addition of cytokines blockers (Figure 11C, D, G, H, J, L, N & P) as reported and evaluated by other studies (7,10,15). This data (Figure 11) supports that despite the cytokine blocker specific downregulation of 4-1BB, it is possible to overcome and capture the magnitude of stimulant-specific differences in a clinical cohort by ICS or nuclear stain. Therefore, researchers should be aware of sensitive markers like CXCR5 and 4-1BB and should validate their usage if they wish to change the stimulant other than mitogen or peptides, especially if they run ICS or nuclear staining. In the final panel, nuclear staining was performed as the primary objective, as this panel will be utilized in the OUD cohort, where FoxP3^+^IFNγ^+^ cells play a pivotal role (4). Notably, no protocol-specific (ICS vs. nuclear staining) differences in the AIM profile were observed (Figure 11I-P).

**Figure 11:**
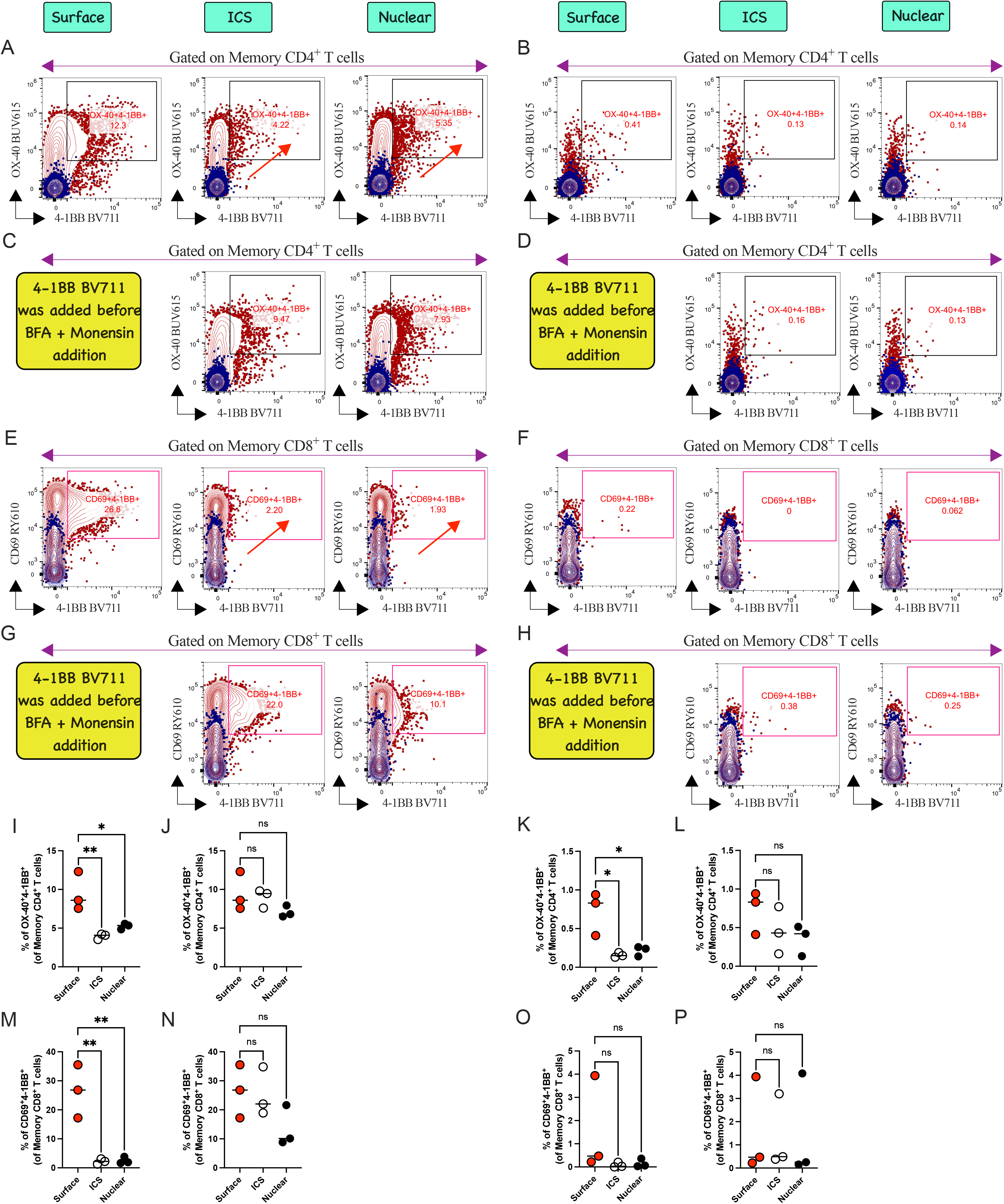
Validation of CD137 (4-1BB) expression upon Con A (A, C, E & G) or CEFSX Ultra SuperStim peptide (B, D, F & H) stimulation after surface stain, ICS, and nuclear stain. Each column represents the different steps of immunophenotyping as indicated on the top. Activation profiles were visually captured by overlaying blue subsets (steady state) on the brown subsets (Con A or peptide stim, 20 h) in the representative bivariate plots (from an individual donor). Graphical presentation of AIM profile after Con A (I, J, M & N) or peptide stimulation (K, L, O & P) illustrates the magnitude of the kinetics of the stimulant-specific 4-1BB profile either in memory CD4^+^ or CD8^+^ T cell compartments. 4-1BB expression after Con-A stimulation was significantly downregulated (red arrow) during traditional ICS and nuclear stain (A, E, I, M). Such significant downregulation can be easily overcome by adding 4-1BB (as indicated in yellow boxes) before the cytokine blocking step (C, G, J, N). Such a pattern was re-evaluated using (K, L, O & P) peptide stimulations. Statistical comparison was performed using either 1-way ANOVA or nonparametric Kruskal–Wallis test corrected for multiple comparisons; *P <0.05, **P <0.01, ns denoted non-significant. Each circle represents a single participant (n=3 per group).

##### 3.3.2.3| Differences in Cytokine Profile between ICS and Nuclear Stain and Kinetics

We validated the effect of ICS or nuclear stain or protocol specific changes in rare Th1/2/17 cells compartment after Con A or peptide stimulation. We believe that head-to-head validation is necessary to understand protocol-specific alterations of rare Th signatures, if they exist. We did not find such details in previously published OMIPs, prompting us to conduct a comprehensive analysis. Our analysis is limited to AIMs (Figure 11) and Th polarization, as well as Treg signatures (Figure 12), as these were the primary focus of our research.

**Figure 12:**
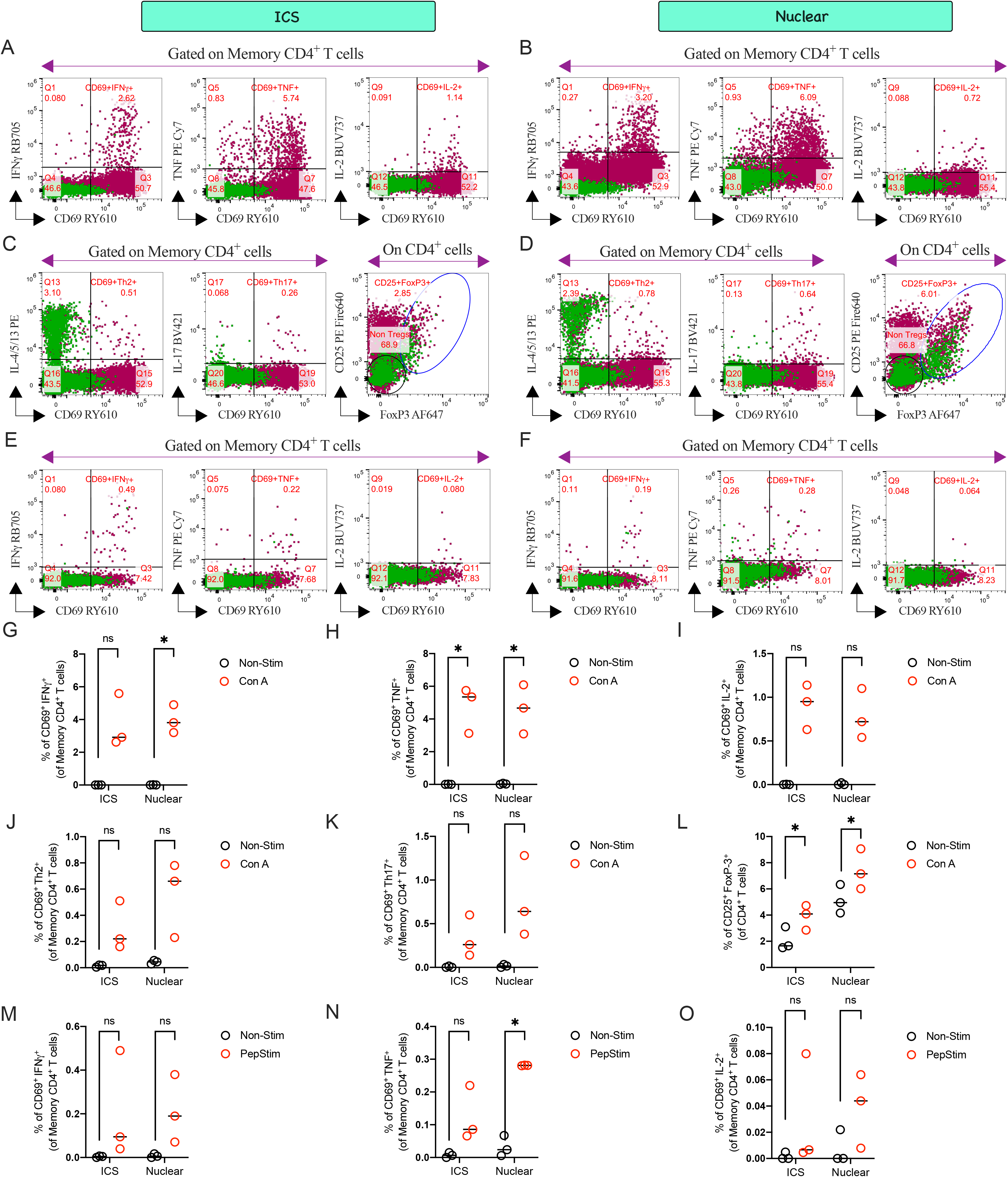
Immune-profiling of cytokines and FoxP3 signatures in stimulated conditions using ICS or nuclear stain. (A, B) Intracellular Th1, (C, D) Th2, Th17, and FoxP3 profiles were compared with either ICS (A, C) or nuclear stain (B, D) after Con A stimulation. CEFSX Ultra SuperStim peptide specific Th1 responses were also evaluated using (E) ICS and (F) nuclear stain. Activation profiles were visually captured by overlaying clover (steady state) subsets on cayenne subsets (Con A or peptide stim, 20 h) in the representative bivariate plots (from an individual donor). (G-K) Graphical presentations of Th1/2/17 profiles after Con A stimulation either via ICS or nuclear staining. Both staining procedures effectively captured the Th1 profile in memory CD4^+^ T cells upon Con A stimulation. Nuclear staining demonstrated superior efficiency in capturing the (G) IFN*γ* signature. As anticipated, nuclear staining (D & L) exhibited excellent efficiency in targeting FoxP3^+^ Treg cells within the CD4^+^ T cell compartment. Conversely, ICS resulted in (C & L) suboptimal capture of the FoxP3^+^ Treg signature. Finally, our panel has been validated to target (M-O) peptide-specific Th1 responses, and (N) significant TNF responses were successfully captured using nuclear staining. Statistical comparison was performed using either two-tailed paired t-test or Wilcoxon test corrected for multiple comparisons; *P <0.05, ns denoted non-significant. Each circle represents a single participant (n=3 per group).

In human, we were unable to capture any protocol-specific differences in the Th1/2/17 compartment except for a better resolution of Con A-specific IFNγ responses (Figure 12G) and peptide-specific TNF responses (Figure 12N) during nuclear staining. Subsequently, we captured the FoxP3^+^ Treg signature using conventional ICS (Figure 12C & L) as well as the recommended nuclear staining protocol (Figure 12D & L). Unexpectedly, we observed that both staining protocols could capture the nuclear FoxP3^+^ Treg signature (Figure 12L) in both steady-state and upregulated states upon Con A stimulation. We recommend following the conventional nuclear staining protocol to capture the Treg signature, as it preserves the resolution and separation of CD4^+^CD25^+^FoxP3^+^ Tregs better during nuclear staining (Figure 12C, D & L).

As we captured delayed cytokine responses at 20 h, therefore we further compared its profile with early cytokine release at 6 h in adaptive T cell compartment (Figure 13). We did not capture any significant differences in adaptive cytokine profiles, but both ICS (Figure 13F) and nuclear staining (Figure 13L) provide better resolution during early TNF detections. Of note, we validated total cytokines on Memory CD4^+^ or CD8^+^ T cells as 6 h stimulation does not induce AIM profile of CD69 successfully (Figure 9K). After 20 h of stimulation, in the final panel, we profiled adaptive cytokine responses as CD69^+^ Th^+^ cells (Figure 1E, Figures 9N, Q, T & Figure 12) as per OMIPs-5(18), 52(19), 67(20) and 91(21). This data supports the effectiveness of both ICS and nuclear staining in capturing late Th signatures using our 28-color spectral panel.

**Figure 13:**
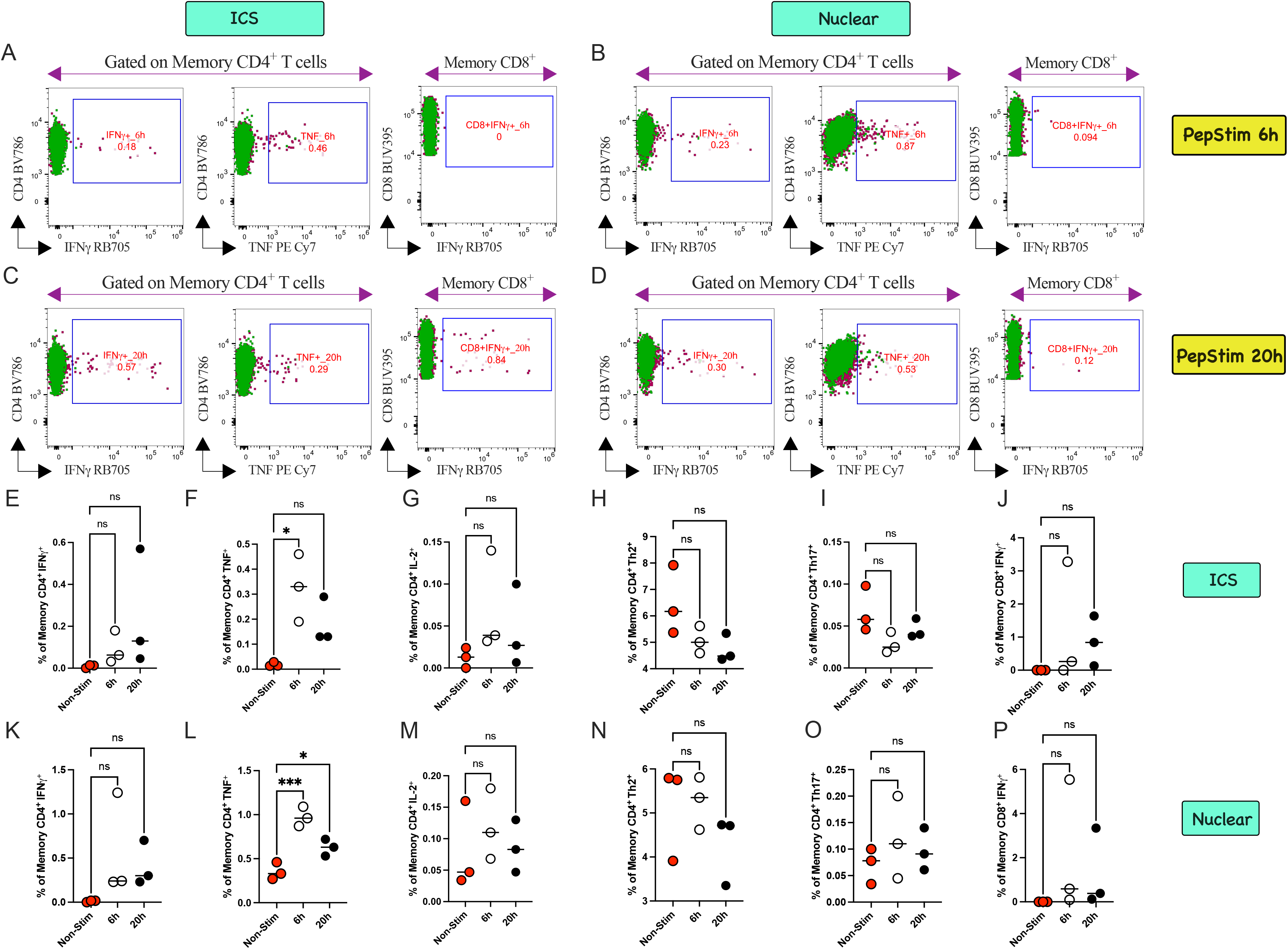
Kinetics of cytokines profile in adaptive immune compartment after peptide stimulation. Bivariate plots comparing early (A, B) and late (C, D) adaptive cytokine profiles either by ICS (A, C) or nuclear stain (B, D). Graphical presentation of ICS profile (E-J) and nuclear staining profile (K-P) illustrates the magnitude of the kinetics of the antigen-specific Th1/2/17 and CD8^+^IFN*γ*^+^ profiles. After resting, cryopreserved PBMCs from healthy human donors (N=3) were stimulated with the CEFSX Ultra SuperStim peptide pool for either 6 h or 20 h. Cytokines were blocked using BFA and monensin for the final 5 h. Statistical comparisons were conducted using either a one-way ANOVA or a nonparametric Kruskal–Wallis test, corrected for multiple comparisons. *P <0.05, ***P <0.001, and ns denotes non-significant. Each circle represents a single participant (n=3 per group).

#### 3.3.3| Data Analysis

Data were collected using a Sony ID7000^TM^ Spectral Cell Analyzer (see **Section 2.1**) equipped with five lasers at the Cytometry Cores of DFCI in Boston, MA. After running single color reference beads/cells and defining unmixing, test samples were analyzed. For the full panel run, a stop gate was activated on live CD45^+^ cells with a cell count of 110K. Upon the completion of the acquisition, all fluorochrome combinations (N x N plots) were meticulously analyzed using the “unmixing viewer”. The quality assurance of all fluorochrome combinations (N x N plots) was further conducted by a Flow Cytometry Technician at DFCI Cytometry Cores to verify the signal intensity of each marker during data acquisition and its distribution in the N x N plots of the “unmixing viewer”. No manual adjustments were made to the SSM. Notably, no unmixing issues were observed during the experiment, as a combination of beads and cells was utilized to define the SSM (Figure 8).

Generated FCS files were analyzed using FlowJo 10.10.1 for Mac Operating System. The gating strategy for the final panel after nuclear stain using human PBMCs was demonstrated in Figure 1. The NHP counterpart of this panel was subsequently validated using PBMCs from AGMs and baboons (1). Figures 11 and 12 demonstrated the consistency of the panel in detecting TCR-independent or dependent AIMs and Th signatures after mitogen or peptide stimulation, respectively. High-dimensionality (HD) reduction analysis was conducted utilizing the Fast interpolation-based t-SNE (FIt-SNE) algorithm on the OMIQ platform (Table 5). For the FIt-SNE algorithm run, we adopted the parameters from our previous study (65). The OMIQ platform sampled 110K events of live CD45^+^ populations equally per file using the “Subsampling” task. We included features (fluorochromes) that specifically targeted the desired cell population, enabling the visualization of those cells within the cellular landscape (Figure 1G). For each FIt-SNE algorithm run, the parameters were set as follows: Number of dimensions = 2, Perplexity = 50, Theta = 0.5, Random Seed = 7000, Max iterations = 1500, Stop Early Exaggeration = 250, Learning Rate = 5000 and Early Exaggeration Factor = 12. By FIt-SNE embedding, we overlaid manually gated major innate and adaptive immune cell lineages in the cellular landscape of live CD45^+^ subset (Figure 1G) for concatenated visualization either in steady state or in Con A stimulated conditions.

## 4.| Data Availability Statement

Data are available in the main article, figures, and tables. Fully stained, quality-controlled, and assured spectral flow data (FCS files) can be accessed through the online repository platform Zenodo (https://doi.org/10.5281/zenodo.23605896). The deposited data will be made publicly available upon publication. The repository contains compressed (.zip) folders of unmixed FCS files for single-color controls and fully stained human PBMCs (stimulated and unstimulated), alongside a standalone FlowJo workspace (.wsp) file illustrating the gating strategy. Each folder includes a “Sample Description” PDF-generated from workspace screenshots, mapping individual FCS filenames to their corresponding sample conditions. Additional data that support the optimization of the study (i.e., Titrations, FMOs, related to Figures 9-13, particularly Surface and ICS profile) are available from the corresponding authors upon reasonable request.

Taken together, our meticulously optimized panel distinguishes itself by its exceptional ability to simultaneously capture antigen-specific AIMs, intranuclear Th profiles, and nuclear Treg signatures.

## Author Contributions

**Soumik Barman:** conceptualization, methodology, data curation, investigation, validation, formal analysis, visualization, writing-original draft, writing-review and editing.

**Arsh Patel:** conceptualization, software, validation, formal analysis, visualization, data curation, writing-review and editing.

**Aisling Kelly:** methodology, writing-review and editing.

**Ben Janoschek:** conceptualization, software, validation, formal analysis, visualization, writing-review and editing.

**Ravi Hingorani:** conceptualization, validation, writing-review and editing.

**David J. Dowling:** conceptualization, supervision, funding acquisition, project administration, resources, writing-review and editing.

## Acknowledgements

The *Precision Vaccines Program* (PVP) is partially supported by Boston Children’s Hospital, Department of Pediatrics. The authors express their gratitude to the members of the PVP and the Program Director, Dr. Ofer Levy, for their valuable discussions and support. Additionally, the authors acknowledge the support and feedback provided by Suzan Lazo (Director, Cytometry Cores, DFCI) and Matthew Powers (Assistant Director, Cytometry Cores, DFCI). Furthermore, the authors extend their appreciation to the late Dr. Andrew Draghi II from Sony Biotechnology Inc. for his insightful input regarding the Sony ID7000^TM^ Spectral Cell Analyzer. Soumik Barman extends his sincere gratitude to Dr. Rajesh Panigrahi of Waters Biosciences for his invaluable guidance related to Real Blue fluorochromes during the panel optimization. The authors also thank Eric Haas (Ionic Cytometry Solutions) for assistance with quality assurance of the deposited data.

## Supporting Information

Additional supporting information can be found online in the Supporting Information section. **Data S1:** Cytometry Part A Author Checklist: MIFlowCyt-Compliant Items.

